# Twainspotting: Identity Revealed During a Simple, but Extended Conversation with a Humpback Whale

**DOI:** 10.1101/2023.02.12.528197

**Authors:** James P. Crutchfield, Alexandra M. Jurgens

## Abstract

Extended acoustic interactions with a humpback whale (*Megaptera novaeangliae*) were captured via human-initiated playbacks of the purported humpback “throp” social call and hydrophone recordings of the animal’s vocalized responses during August 2021 in Frederick Sound, Southeast Alaska. Multivariate statistical analyses performed after the event, that adapted the Wasserstein metric to spectrograms, strongly imply that the played back call was from the same animal, which had been recorded the previous day and initially selected for its clarity. This suggests these interactive playback studies may have formed a probe of time-delayed self-recognition. Fluke photographs taken that previous day and during the interactions revealed that the animal had been identified 38 years earlier, now known as the female humpback named Twain 14 years ago.

This exciting and to our knowledge unique series of interactive vocalization events, while hopeful, poses more questions than answers. Most basically, did the half-hour long series of acoustic exchanges constitute an interspecies conversation? We argue that analysis tools available to infer the implied causality of interaction—statistical dependency, multivariate information theory, and machine learning—leave the question open. That said, and perhaps more importantly, the extended interaction broaches questions whose answers bear directly on future interspecies communication and, more generally, the human appreciation of nonhuman intelligence. The reported human-humpback interactions will facilitate exploring these issues at new depths.

---

… there may indeed be no possibility of finding out the answer … but … grateful for the crumbs of truth—the minor insights—that whales deign to drop from time to time in front of scientists like me.

Roger Payne, *Among Whales* [1, pg. 111].

## I. INTRODUCTION

First appearing in the Earth’s oceans some 40 Myrs ago, cetaceans exhibit compelling evidence for advanced intentional behaviors and conscious awareness through their raw intelligence, song generation [2, 3] and sharing [4, 5], communication and interactions with their own and other species [6, 7], and empathy (concern for others’ well-being) [8]. Over this long evolution—exceeding humans’ by a factor of 10—they developed tools (socially-coordinated bubble-net feeding by humpbacks) and region-(and possibly hemisphere-) spanning ocean-acoustic communication networks [9]. Over the last half century humpback whales, in particular, became known for their active vocalizations. These fall into two categories: One *comprised of extended songs* (minutes to hours), emitted predominantly by males; the other *social calls*, short vocalizations (lasting seconds) that occur in animal interactions and are produced by both males and females [2].

The cetacean world-experience—more succinctly, a whale’s *Umwelt* [10]—is predominantly acoustic. Water visibility is low with high turbidity and little light at typical diving depths (> 50m). Though surfacing regularly to breathe, whales spend the bulk of their time below the ocean surface, often for extended periods. These are the challenging circumstances that science must meet to study whales.

Given a watery world, the experience of which is mediated by sound, how does whale cognition manifest in their communicative acoustic interactions? Success in addressing this would not only substantially enhance their conservation, but also advance our appreciation of co-existing and independently intelligent animals on earth. The research challenge is substantial, though, since properly meeting them one must study these animals acoustically via hydrophones only occasionally supplemented with surface visual and acoustic observations, and even more rarely digital video-sound recording tags [11].

The following recounts an event bearing directly on these concerns: Extended acoustic interactions with a humpback whale (*Megaptera novaeangliae*) during August 2021 in Frederick Sound, Southeast Alaska. Over a near half-hour period, the interactions consisted of a series of (i) acoustic playbacks (broadcast via underwater loudspeaker) of the purported humpback “throp” social call [12] spaced (contingent on response) approximately a minute apart, along with (ii) hydrophone recordings of the animal’s immediate throp response vocalizations and (iii) visual and photographic tracking. The sequence of human-initiated playback and animal response was almost pure turn-taking [13].

The interaction in question occurred midday in Frederick Sound during a week-long circumnavigation of Admiralty Island in southeast Alaska aboard R/V Glacial Seal (Juneau, AK). This location is known for its humpback whales who visit during their summer feeding season (June-September) and then winter over in Hawai’i for breeding and calving (December-March). Figure 1 shows the field research configuration with vessel and electronics, including hydrophone, underwater loudspeaker, and animal in relative positions.

**FIG. 1:**
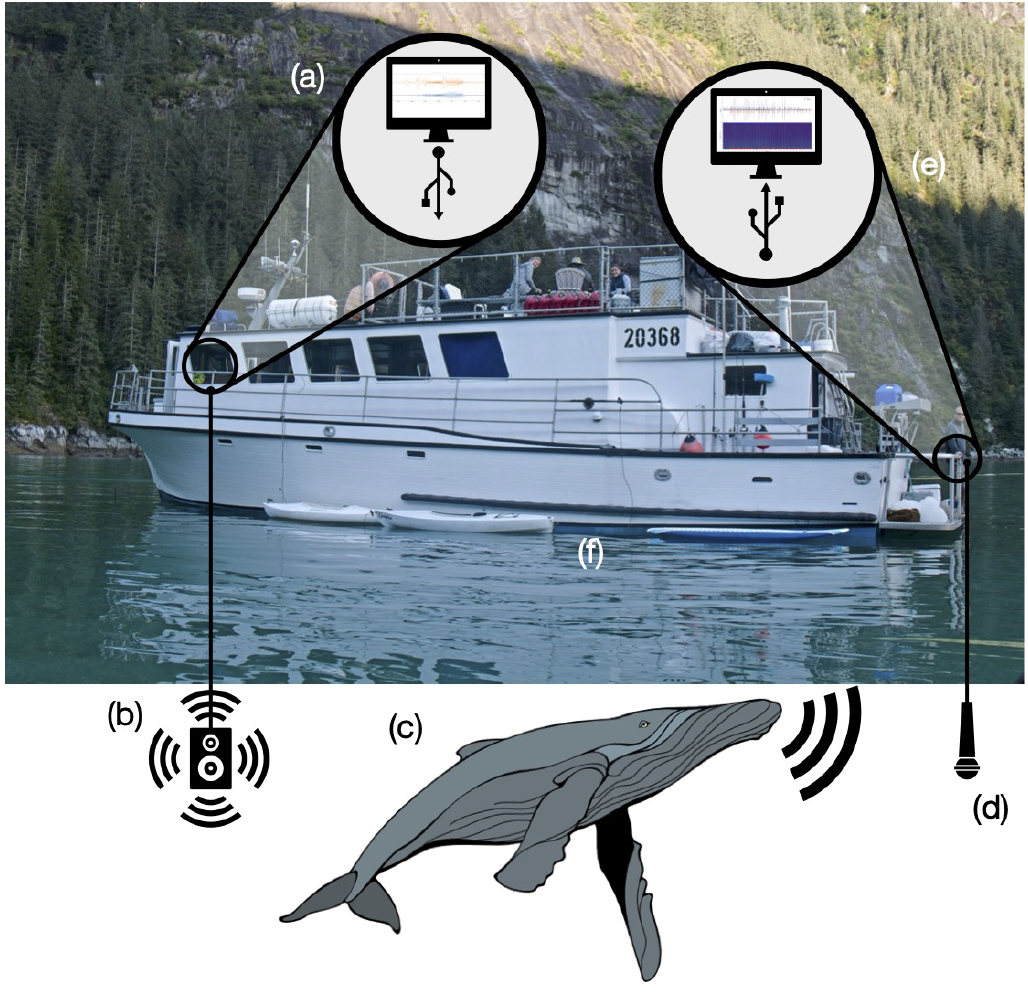
Interactive communication in the field: (a) loud-speaker playback computer (MacBook Pro 15”), (b) underwater loudspeaker (Lubell LL916C-025’) submerged driven by audio amplifier (Lubell modified TOA CA160), (c) humpback whale (*Megaptera novaeangliae*), (d) hydrophone (C54, Cetacean Research Technologies), (e) hydrophone recording computer (MacBook Pro 15”), and (f) vessel (R/V Glacial Seal, Juneau, AK). Video recordings and photographs made by observers on top deck; computers with human operators inside on main deck.

Specifically, we report on a series of 36 repeated playbacks and subsequent animal throp vocal responses. To test the causal interdependency of interactive two-party communication, the operators on the main deck terminated playbacks after about 20 minutes and the animal responded in a delayed fashion—an immediate single throp call after a quarter minute, then another two each separated by 42s, and then a final call 41s later. The final calls occurred with diminishing acoustic power, as the animal swam away (verified visually and photographically). No further throp calls were heard. The several nearby humpbacks (< 1000m) did not vocalize during the acoustic exchanges.

Notably, the actual playback throp call used was captured the previous day in approximately the same location. It was employed during the next day’s interactive playback trials due to the hydrophone recording’s high acoustic quality—low sea-state and ambient noise and temporal isolation from other animal vocalizations.

Shortly after the interaction an online [humpbackfluke database](https://happywhale.com/home/) identified the animal from fluke photographs (Figs. 4a and 4b) taken during the interaction and the previous day’s sightings as one and the same—a female humpback (ID SEAK-0401) first identified 38 years earlier in southeast Alaska [14]. She was sighted a dozen years ago by Fred Sharpe (Alaska Whale Foundation) who named her Twain at that time [15]; see Fig. 4c.

In addition to this photographic evidence, analysis of the spectra of the playback and response recordings reveal that the playback call and the responses are not distinguishable. Taken together with corroborating inferences from contemporaneous visual sightings, one concludes that the call used for playback was the same animal as the one present during the interaction. Thus, the interactions appear to be a *time-delayed vocal self-recognition study*. Self-recognition studies are an experimental protocol familiar from behavioral studies of other animals in which mirrors are deployed for visual recognition. While visual examples been practiced with dolphins [16], we believe this example of a delayed vocal self-recognition to be unique for whales.

To interpret these observations, the following first provides background on the relevant marine biology of cetaceans and humpback whales emphasizing, in particular, vocalization, communication, and behavior studies. This review highlights that Twain’s behavior was highly unusual, especially when compared with prior behavioral response studies (BRS) with mysticetes.

With the setting laid out, it turns to describe the acoustic interactions and statistical results, outlining how the analyses answer several questions related to animal identity and human-animal communication. It then briefly comments on the challenges of quantitatively establishing the existence and performance of interactive communication using multivariate statistical, information-theoretic, and machine learning methods. It closes with a discussion of possible interpretations of the field observations and quantitative analyses, including extant statistical methods’ substantial limitations. The conclusion leverages these to present several forward-looking challenges to future investigations of nonverbal animal and interspecies communication.

Supplementary Materials (SM) provide a summary of the natural history of humpback whales, the history of acoustic monitoring of cetaceans, details of our fieldresearch methods, theoretical detail on statistical signal analysis and information-theoretic tools, hydrophone recordings of playbacks and animal responses, associated waveforms and spectrograms, and a narrative timeline of interacting with Twain.

## II. HUMPBACK NATURAL AND RECORDED HISTORY

Fairly interpreting the extended interactions with Twain requires appreciating the natural history of humpback whales and the long history of underwater acoustic monitoring. The following gives brief summary of relevant factors that are detailed in SM A.

The product of evolution over 25 Myrs, today’s mysticete (baleen) species of humpback whales (*Megaptera novaeangliae*) are found in all the planet’s oceans. They migrate annually between high-latitude coastal summer feeding grounds and low-latitude shallow-water breeding and calving grounds. Of the North Pacific population, a significant fraction winters each year in waters surrounding the major Hawaiian islands and returns to Alaskan waters during the summer months to feed [17–19]. The social behavior of humpback whales while in Hawaiian waters is largely related to reproduction. Humpbacks are one of the most vocal whale species, which is particularly evidenced during mating interactions.

Considering that cetaceans function wholly in water, a dense medium in which light attenuates far faster than sound, it is not surprising that hearing and vocalizing are believed to be their fundamental sensory and communication channels. Their infrasound vocalizations are implicated in maintaining the stability of their herds in typically low-visibility ocean waters [20]. Cetaceans have a 10-11 octave functional hearing range compared to human’s 8-9 octave range and a maximal functional high frequency capacity of 20-30 kHz. The consensus, though, is that humpback (generally mysticete) ears are adapted for good low to infrasonic hearing.

Given their water milieu and activities that keep them below the surface for extended periods of time, acoustic monitoring of humpbacks is a primary method of investigation [21]. There is a long history of passive acoustic monitoring and acoustic playback studies going back to the 1950s; though sea-faring peoples have heard them for millennia through the hulls of their watercraft [22]. Playback studies showed, overall, that whales respond to recordings of their own sounds in complex, context-dependent ways. For example, both syntax (combining individual sounds into a series) and context were important in conveying information via playbacks [23].

Most dramatically, adult humpbacks exposed to social sounds approach playback vessels [24]: it is “impressive to an observer in a 5 m boat when a 15 m whale charges the boat.” The whales that charged swam at the surface approaching to less than 5 m from the underwater playback speaker and then “swam around the boat in ever widening circles, occasionally making another pass under it. Although charging whales came as close as 2 m to the speaker, they never touched it”.

The acoustic interactions reported used the humpback “throp” call (sometimes described as “whup”). Purportedly, this is a social greeting call [12]. The vocalization starts out as a low-frequency amplitude-modulated call (“growl”) that ends with a increasing-frequency sweep. Humpback whales across all oceans vocalize throps. The call has persist across generations. The throp call is believed to be innate in this species [25–29].

Given the long history of playback studies, several contrasts with the present approach and results require highlighting. In short, the interactions with Twain are atypical in light of previous reports of humpback social vocalizations. First, previously social sounds occurred largely in groups containing three or more whales. Social sounds were infrequently heard near single whales, pairs, or cow-calf groups. Our playback interactions involved only one vocalizing animal—Twain. Others were present, but did not vocalize; see Fig. 3. Second, the interaction occurred in humpback summer feeding grounds—southeast Alaska, not as previously in their Hawai’i winter calving grounds. Third, in the summer feeding grounds little to no song is vocalized and little aggression is observed, likely due to the absence there of reproduction activity and mating competition. Fourth, previously social sounds were heard rarely in nonaggressive situations. In contrast, the Twain interactions took place in a nonaggressive setting. Fifth, previously interactions were reported to fall into two exclusive classes: (i) rapid approach to the broadcast vessel or (ii) nonapproach (move-away from vessel). Twain stayed near the vessel for a half hour. Finally, and perhaps most importantly, previous studies did not employ mutual acoustic exchanges. Certainly, what communication there was in previous reports was not interactive, in contrast to the repeated interactions with Twain.

## III. VOCAL INTERACTIONS

This sets the background necessary to appreciate interacting with humpback whale Twain. As we will note, though the field work was undertaken with the full intention to acoustically interact with humpbacks, the particular circumstance reported here was in many ways unintended and certainly surprising.

From the visual sightings, photographs, video recordings, and acoustic analyses of the hydrophone recordings one concludes that the interactions engaged a single animal—humpback Twain. The events of most interest consist of extended, sequential vocal interactions with Twain on 19 August 2021. Over a half-hour period, the interactions consisted of a series of (i) human-initiated acoustic playbacks (broadcast via underwater loudspeaker) of the purported humpback throp social call [12]—the *exemplar* —spaced (contingent on response) approximately less than a minute apart, along with (ii) hydrophone recordings of the animal’s immediate throp-response vocalizations. Figure 2 shows both the hydrophone waveform and spectrogram during the entire interaction, along with annotations of exemplar playbacks (downward blue or white triangles, *P*_0_ − *P*_35_) and Twain’s response vocalizations (red upward triangles, *R*_0_ − *R*_35_).

**FIG. 2:**
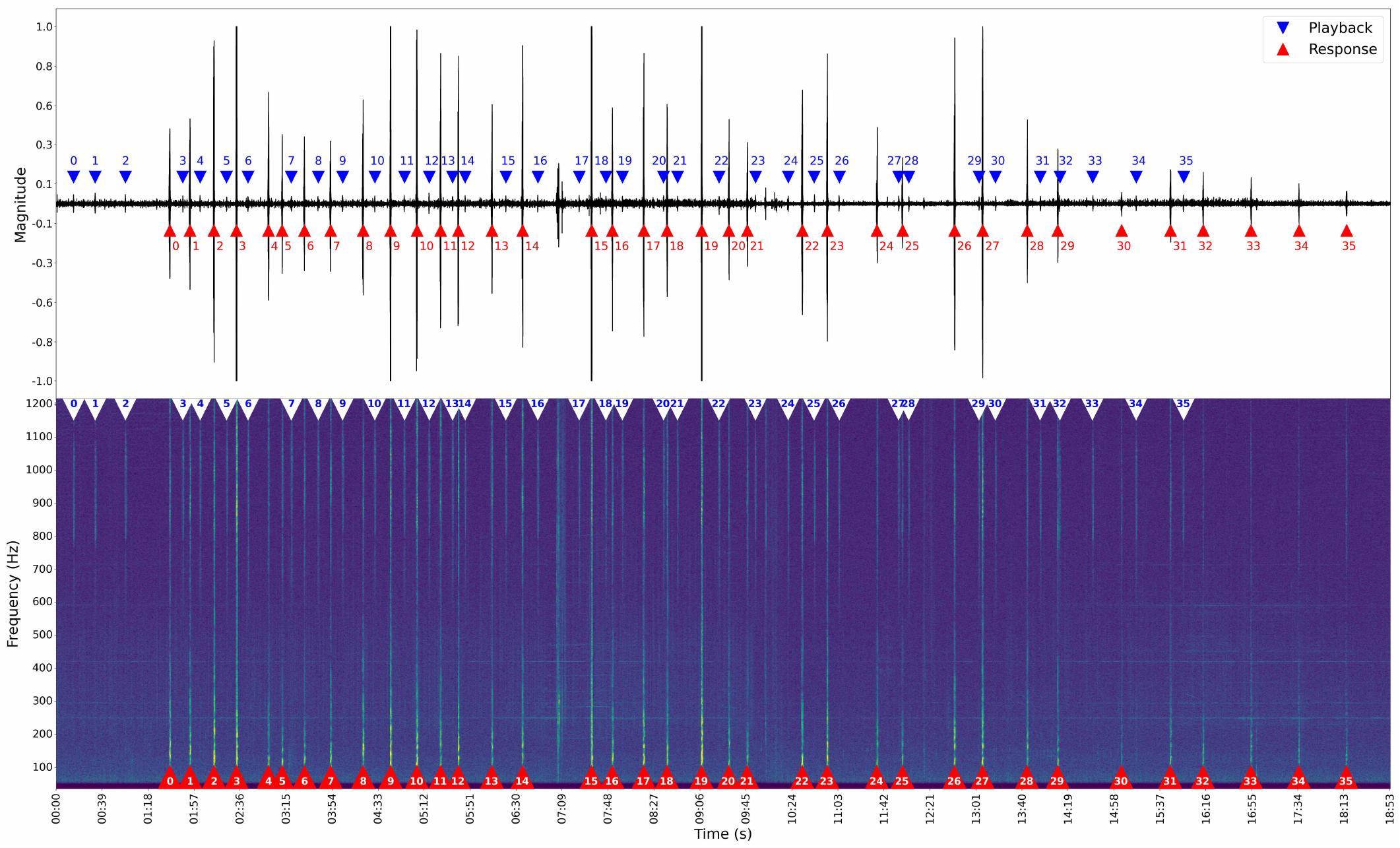
(Top) Waveform of entire acoustic interaction; 19-minute hydrophone recording. (Bottom) Spectrogram for same. Animal responses are clearly identified both in energetic spectrograms and in waveform plots as large-amplitude spikes (red upward triangles). The human-initiated playbacks, however, being markedly lower amplitude and acoustically less powerful, appear in the spectrogram as small spikes (blue or white downward triangles) interpolated between the large (animal-response) spikes. Hydrophone recording 20^th^-order Chebyshev II bandpass filtered to [40, 5000] Hz; 44.1 kHz sampling frequency.

During much of the vocal interaction, the whale was underwater. Figure 3 displays the animal’s surface track near the vessel during the interactions, including nearby animals that did not acoustically interact. The animal surfaced six times (1 − 6 in Fig. 3), some with multiple respirations per surfacing. Based on surface timing and orientation, the whale traveled around the vessel moving port to starboard within a broad half circle at ≈100 m [30]. Before terminating playbacks, the whale doubled back (11:31 AM) and, then, departed to the north (11:34 AM). Once the last playback (*P*_35_) was broadcast the animal subsequently vocalized four additional times (*R*_32_ − *R*_35_).

**FIG. 3:**
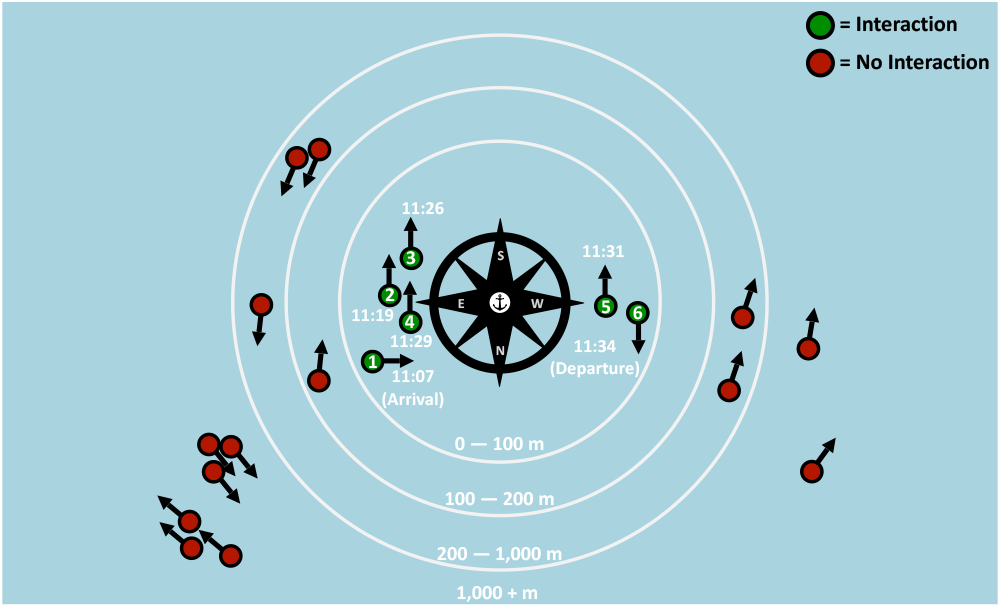
Whale group during Twain’s extended interaction: (Inner circle, 100m) Twain’s surfacings (green circle locations of animal at surface, black arrow orientations) and times. Other whales, in the nearby group denoted with red circles, did not acoustically interact. Adapted from Ref. [30] with permission.

Altogether there was a series of 36 repeated playbacks and subsequent animal throp vocal responses. See the inter-communication timings in Table S1. The sequence of human-initiated playback and animal response was nearly pure turn-taking [13]. Aside from the initial playbacks and final animal responses, the exceptions occurred at responses [*R*_4_, *R*_5_] and at playbacks [*P*_16_, *P*_17_].

Supplementary Material C describes the first interaction with the (as-of-then unidentified) animal on 18 August and Fig. 5 (Top) gives the spectrogram of hydrophone recording of the throp call selected for use the next day. Humpback Twain’s locations and times and those of other nearby humpbacks during the interaction are annotated on Fig. 3 and described in SM D.

The extended interaction occurred the next day (19 August) in approximately the same location as the previous day in Frederick Sound. SM D describes the overall interaction, presents spectrograms of animal physical sounds and vocalizations, playback-response (hydrophone recording) spectrograms and video, animal surface locations and timings, and fluke photographs for animal identification. Links are provided to the online WAV file**1** of the hydrophone recording shown in Fig. 2 and to the video recording with time-aligned hydrophone recording.

A key question, then, is what kind of vocal interaction occurred? To test the causal interdependency of interactive two-party communication, we intentionally terminated playbacks (*P*_35_) after about twenty minutes. The animal responded in a delayed fashion—*R*_32_ − *R*_35_: an immediate single throp call after a quarter minute, then another two each separated by 42s, and then a final call 41s later. The final calls occurred with diminishing acoustic amplitude, as the animal swam away (verified visually and photographically). No further throp calls were heard subsequently.Notably, the several nearby humpbacks did not vocalize during our interacting with Twain. These other humpbacks and their approximate distances to the vessel at various points during the interaction are annotated on Fig. 3.

SM D3 describes the composite recording that merges the camcorder video and audio taken from the top (observation) desk and the hydrophone recording during the entire extended interaction. It provides a link to the online recording.

## IV. IDENTITY

The extended vocal interaction naturally raises the question of who the animal was and what the interactions could mean, if anything. Key to this are visually and acoustically identifying the animal.

### A. Visual Identification

Figure 4(a) presents the fluke photograph identifying the animal encountered on 18 August. Figure 4(b) is that taken on the 19^th^ at the end of the acoustic interaction. And, for historical interest, Fig. 4(c) is a fluke photograph taken 13 years earlier by Fred Sharpe (Alaska Whale Foundation) who named her Twain at that time. The online whale ID database HappyWhale identified the photographs as the fluke of female humpback SEAK-0401. Reference [14] surveys the recorded history of Twain’s sightings over 38 years in the waters of southeast Alaska and west Maui, Hawai’i, and notes several identifying features on her fluke.

**FIG. 4:**
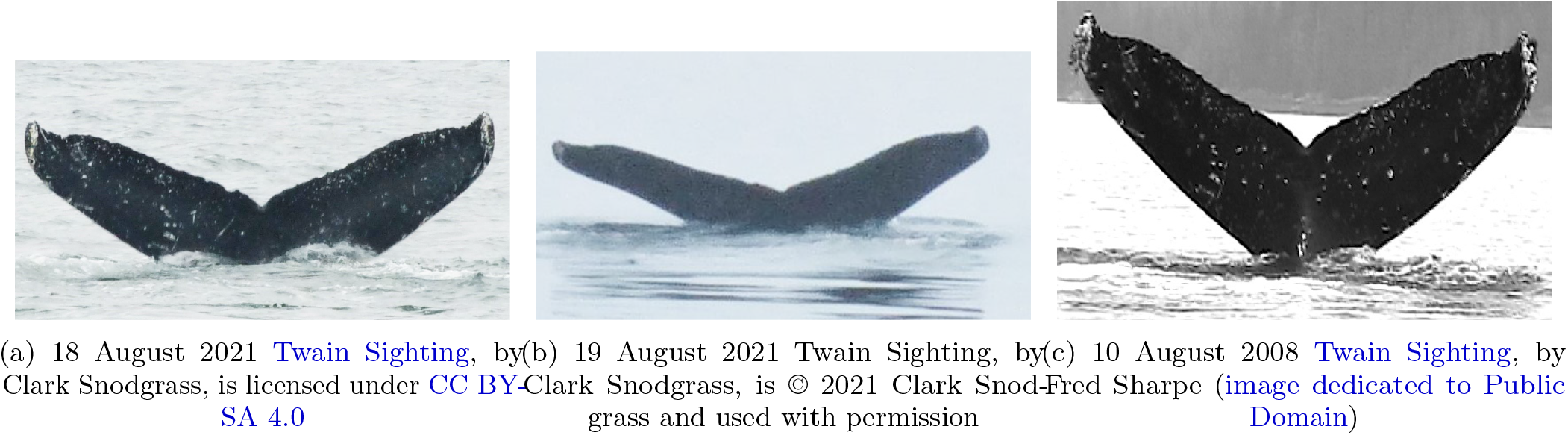
Humpback whale Twain sightings: (a) 18 August 2021, the day the exemplar throp call was recorded. (b) 19 August 2021 end of interactions at point of departure; see Fig. 3 at location 6. (c) Alaska Whale Foundation (AWF) Name: Twain; IDs: SEAK #0401 and HW-MN044056. First Twain sighting was in 1984; most recently seen 22 August 2022 in southeast Alaska; to date 28 total sightings [14].

### B. Acoustic Identity

Along with the fluke photographs and video recording of surface activity, we recorded the animal’s vocalizations via a hydrophone. Figure 5 (Left) shows the spectrogram of the throp call exemplar captured on 18 August. This exemplar was employed for the 19 August interactive playbacks *P*_0_ − *P*_35_ due to its excellent acoustic quality. Figure 5 (Middle Left) similarly shows the first (*R*_0_) and the twentieth (*R*_19_) of the animal’s responses during the extended interactions during playbacks of the exemplar. Figure 5 (Right) gives the response (*R*_35_) spectrogram as Twain swam away, ending the interaction. The similarity of the spectrograms demonstrates that the exemplar and the animal’s response were of the same call type—a throp. These spectrograms and the entire set across the interaction (Fig. S3) demonstrate that the responses were from the same animal.

As Sec. IV A discussed, visual observation combined with fluke identification shots from indicate that the August 18^th^ exemplar throp and the set of thirty-six responses from August 19^th^ were produced by the same animal. To bolster these visual observations we also examine the variation in spectral properties of each sound.

**FIG. 5:**
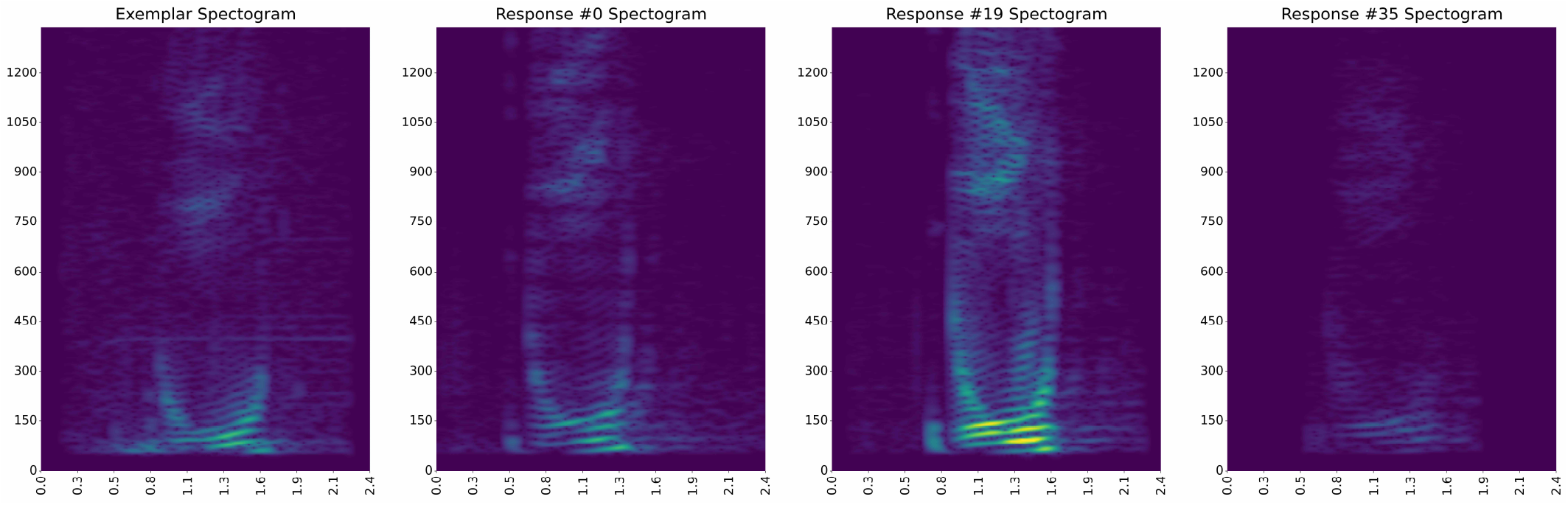
(Left) Spectrogram of Twain’s throp call recorded on 18 August. (Middle Left) Twain’s first response (*R*_0_) to throp playbacks (*P*_0_ − *P*_2_) on 19 August. (Middle Right) Twain’s 20^th^ response (*R*_19_). (Right) Twain’s last (*P*_35_) response as she swam away.

Figure 6 (Top) shows spectrograms of all Twain’s responses (*R*_0_ − *R*_35_) arranged in temporal order. Below, the spectrogram of the exemplar throp sound (*E*_0_) recorded on August 18^th^ is shown alongside each of the August 19^th^ playbacks (*P*_0_ − *P*_35_), again arranged in temporal order. (See Fig. 2 and Table S1 for exact timings and Fig. S1 for details on spectrogram calculation.)

**FIG. 6:**
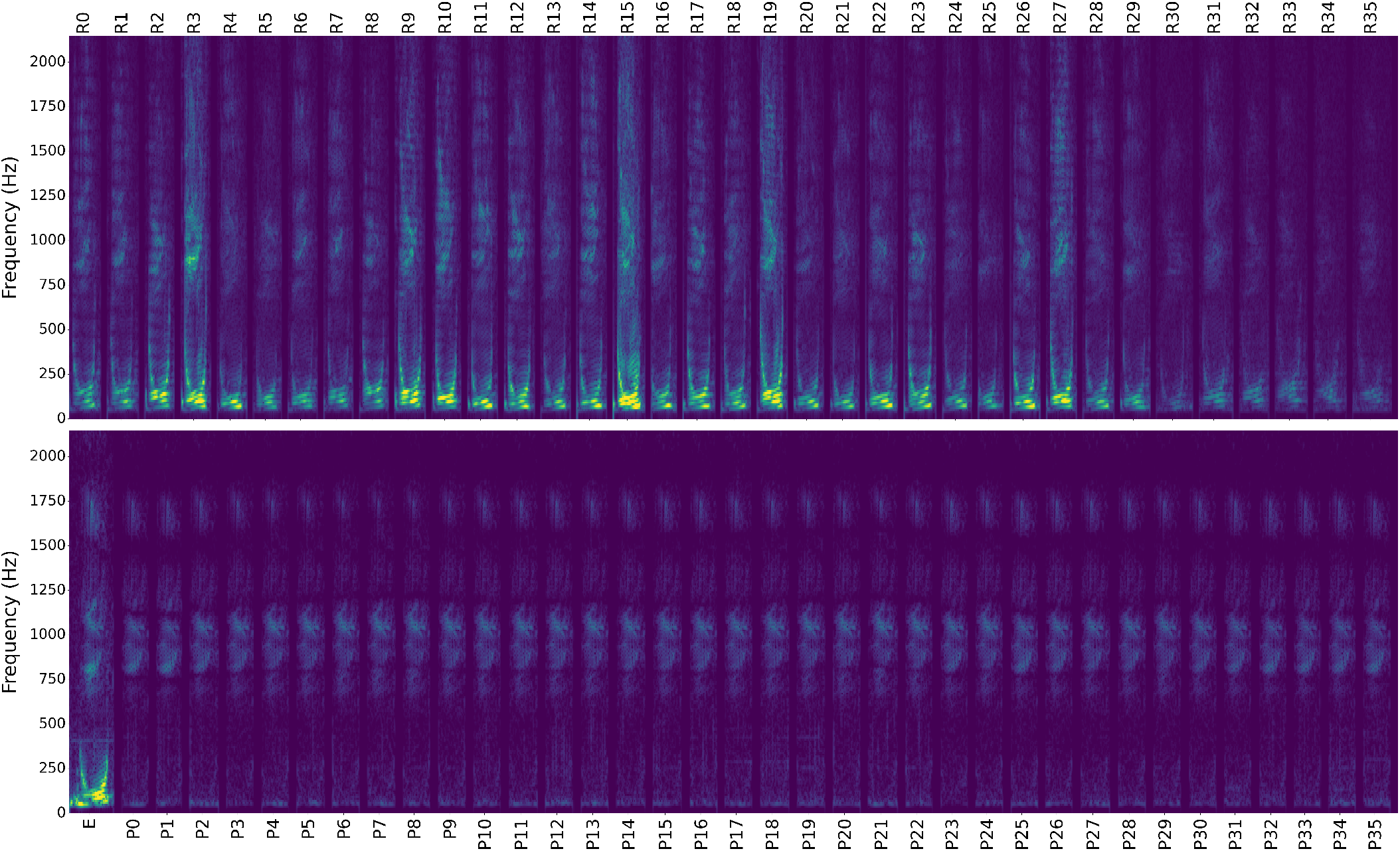
(Top) Spectrograms of Twain’s throp responses *R*_0_ *−R*_35_: Each of Twain’s thirty-six responses clipped out and lined up for simultaneous comparison. Two seconds of silence inserted after each vocalization. Even a cursory visual comparison shows how much variation there is in Twain’s responses, although the characteristic shape of the throp call remains throughout. (Bottom) Spectrograms of the August 18^th^ exemplar *E* and the August 19^th^ playbacks *P*_0_ − *P*_35_. Each playback from August 19^th^ clipped out and lined up for simultaneous comparison with two seconds of silence inserted after each. While the spectrograms of the playbacks do vary according to slight changes in environmental conditions over the course of the interaction, the spectral properties are essentially identical.

The most obvious feature is that the loudspeaker playbacks (*P*_0_ − *P*_35_) of the exemplar throp are anemic in the lower frequencies, completely missing the most characteristic U-shaped band between 40*−* 300 Hz. The higher frequency harmonics are reproduced more or less accurately, although a smooth variation in the frequency spectrum over time can be observed visually across the sequence. As nothing was changed in the equipment or sound being played back, this variation must necessarily be due to shifting ocean conditions. This offers a useful baseline for how much to anticipate recorded sounds vary over the course of the interaction. We also notice that the magnitude of Twain’s responses varied over time, especially for the last three responses when the animal was moving away.

To account for this variation in magnitude, we normalized the energy spectral density of each sound between 40 Hz and 2KHz. This left us with 73 sound spectra—36 playbacks, 36 responses and one exemplar—to compare.

Our chosen distance is the Wasserstein metric calculated pairwise between energy spectral densities. The Wasserstein metric is known as the “earth-mover’s distance” as it estimates how much “mass” must be transported to transform one probability distribution into another. This is useful for normalized sound spectra as they can be thought of as probability distributions of signal energy. Moreover, differences in pure frequencies are weighted by how far apart they are. For eaxmple, the Wasserstein distance between the energy spectra, *µ* and *ν*, of two frequencies, say 440 Hz and 660 Hz, respectively, is simply the difference between the two frequencies: *W*_*p*_(*µ, ν*) = 220 Hz. For more on the Wasserstein metric, see SM F.

Figure 7 (Top) gives the a histogram of the calculated distances, with coloration according to whether the distance is between two playbacks, two responses, or a playback and a response. As noted, differentiating between the playbacks serves as our barometer for how well throp realizations are distinguished. Any variation in sounds below the maximum distance between any two playbacks potentially can be explained by the difference in sea state, recording and broadcasting equipment orientation, and other external changes in experimental conditions.

**FIG. 7:**
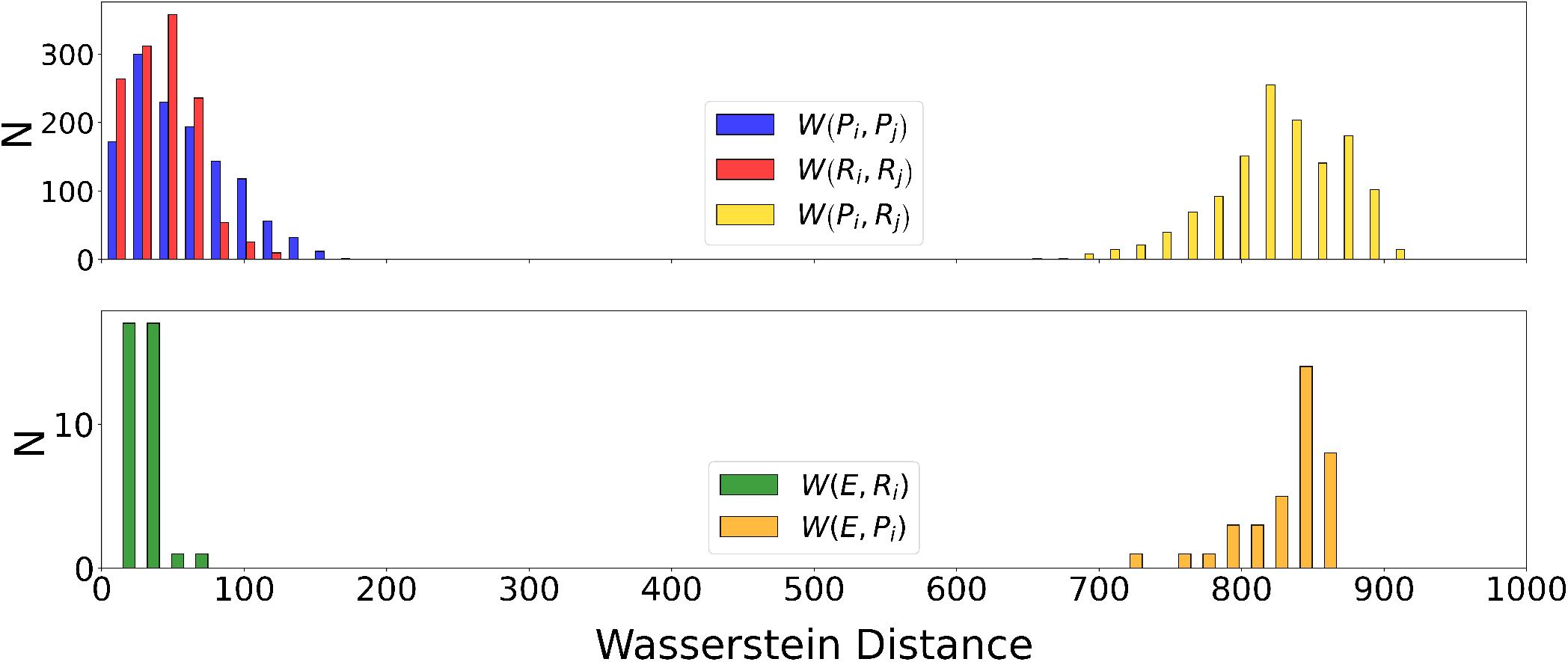
(Top) Histograms of the full-spectrum Wasserstein distances between playbacks *P*_0_− *P*_35_ and responses *R*_0_ *−R*_35_, colored by the distances *W* (·, ·) between two playbacks (*P*_*i*_ and *P*_*j*_, blue), two responses (*R*_*i*_ and *R*_*j*_, red) and a playback (*P*_*i*_) and a response (*R*_*j*_) (yellow). (Bottom) Histogram of the full-spectrum Wasserstein distances between the exemplar *E*_0_ and each playback (*P*_*i*_, gold) and each response (*R*_*i*_, green).

Figure 7 shows that the maximum distance between two playbacks is greater than the maximum distance between any two responses. On the one hand, this indicates that the sounds produced by Twain are remarkably consistent when changes in magnitude are removed. On the other hand, the metric strongly differentiates the playbacks from the responses. This is unsurprising when comparing these results to Fig. 6—illustrating that the most important determiner is the presence-absence of the lower part of the throp spectrum.

Figure 7 (Bottom) shows the distances between the August 18^th^ exemplar *E*_0_ and the playbacks *P*_*i*_ and the responses *R*_*i*_. Similar to above, the exemplar is very close to the responses *R*_*i*_, but well-distinguished from the playbacks *P*_*i*_. The exemplar *E*_0_ falls well within the range of variation for Twain’s throps from August 19^th^.

It is additionally informative to restrict distance calculations to the higher-frequency range—750 Hz to 1500 Hz—to remove the strong biasing produced by the loudspeaker playbacks lacking the throp lower-frequency components. Although this interaction feature is vitally useful information and should not be discarded in our analysis, for the purpose of comparing variation in the frequency spectrum it is useful to compare apples to apples, so to speak.

Figure 8 (Bottom) indicates that when the lower frequency range is filtered out, the difference in variation observed between the exemplar *E*_0_ and the responses *R*_*i*_ and the exemplar *E*_0_ and the playbacks *P*_*i*_ vanishes. In other words, in the higher frequency range, one cannot distinguish between the exemplar, a playback, or one of Twain’s responses. This argues that all of these sounds were created by the same animal.

**FIG. 8:**
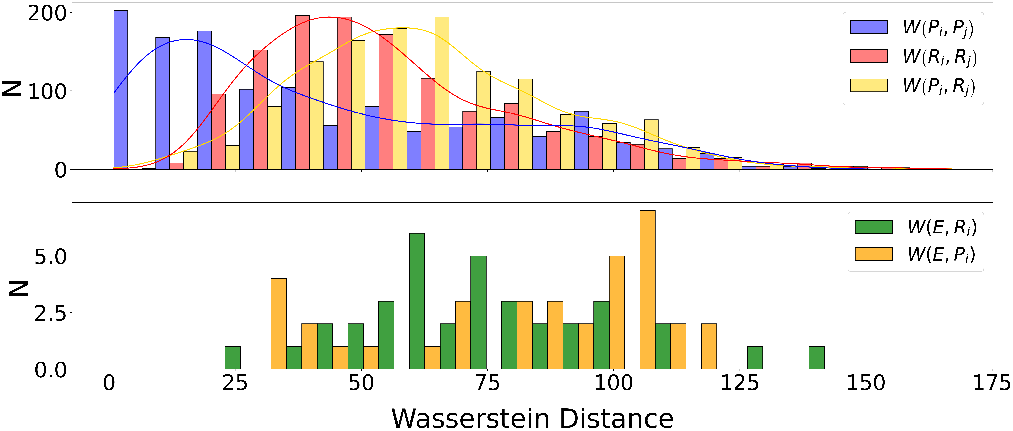
(Top) Histograms of the high-frequency spectra Wasserstein distances *W* (·, ·) between playbacks *P*_0_ *−P*_35_ and responses *R*_0_ *−R*_35_, colored by the distances between two playbacks (blue), two responses (red) and a playback and a response (yellow). Distances calculated only between 750 Hz and 1500 Hz. (Bottom) Histogram of the high-frequency Wasserstein distances between the exemplar *E*0 and each playback (*P*_*i*_, green) and each response (*R*_*i*_, gold). Distances also calculated only between 750 Hz and 1500 Hz.

Thus eliminating the lower frequency components brings the range of distances *W* (*P*_*i*_, *R*_*j*_) between playbacks and responses into line with the variations among playbacks *W* (*P*_*i*_, *P*_*j*_) and among responses *W* (*R*_*i*_, *R*_*j*_). In this case the variation of the responses is on average slightly greater than the variation in the playbacks. Although not a strong effect, it is in line with expectations that the responses vary to a greater degree, even if only due to the fact that repeating exactly the same sound is more difficult for a biological organism than a broadcast digital recording. That said, greater variation in the power spectra’s upper ranges is also intriguing as these ranges may harbor greater semantic information.

These analyses of the vocalizations—exemplar, playback, and responses—lead to the following conclusions:

1. During the extended interaction, hydrophone recordings of the animal’s responses match that from previous day; see Fig. 5;
2. Hydrophone recordings of the animal’s responses during interaction matches that of the exemplar;
3. Spectral analysis of the animal’s responses during the interaction indicate they were very similar; see Figs. 5 and S3. While the statistics are consistent with them being made by same animal, being definitive requires comparison to a range of throp vocalizations from other animals.
4. Spectrogram homogeneity and lack of variation across animal responses during the interaction suggests a lack of semantic variation. However, not knowing at this point humpback communication, at best this is a superficial anthropomorphic observation.
5. Response-time variations, though, might indicate relevant communicative information. See SM D 1, in particular Table S1 and the contingency analyses of SM D 2.

The signal analyses for acoustic matching used energy spectra and spectrograms in several ways. To probe robustness, these analyses were conducted over a range of analysis parameters—sample rate, sliding window size, bandpass filtering, and the like. The SM gives the technical details and further results.

### C. Alternative Analyses

Our wide-ranging statistical explorations of the putative communication exchange leads us to briefly comment on the available quantitative tools. For as long as spectrograms have been readily available, it has been appreciated that they have many weakness, especially when analyzing communicative vocalizations [31, 32]. For a thoroughgoing technical critique of spectral methods used in the service of detecting multivariate statistical dependencies see Ref. [33].

For these reasons, alternative approaches to determining call similarity were also explored. For example, we applied spectral matching [34] and spectral reassignment to extract a template for the throp calls using it to preprocess spectrograms for better matching [35–37]. Ultimately, these statistics were more complicated than and not as insightful as the spectral distances and similarities we report here.

That said, previous uses of communication theory applied to humpback vocalizations [38], led us to pursue information-theoretic measures including time-delayed mutual information, transfer entropy, and related methods [39, 40]. Though their use confirmed the conclusions from the spectrogram analyses, they did not provide markedly improved identification. And so, we do not include their description here, though future analyses may reveal them to be insightful. Finally, and more profoundly, despite their origins in Shannon’s communication theory, information-theoretical methods [41–43] simply do not address the nature of information flow, causality, and spontaneity [39, 40] implied in genuine interactive communication.

### D. Interaction Semantics: Time-delayed self-recognition?

The fortuitous selection of the 18 August throp call and its use during the 19 August extended interactions suggest that during the interactions the animal was hearing broadcasts of her own throp call. And, this suggests the unusual hypothesis that the extended interactions functioned as a kind of time-delayed self-recognition study. Strictly speaking, though, it was not a probe of self-recognition in the sense of previous studies of self-recognition by animals viewing themselves via a mirror [44]; see, in particular, Ref. [16]’s exploring dolphin visual self-recognition. The interaction with Twain involved hearing, not vision. While acoustic mirrors exist [45]—parabolic surfaces that directly reflect sound—this is certainly not what happened physically in the Twain interaction, given the electronic recording and playback. More to the point, the animal’s vocal responses were not immediately played back to the animal.

This all noted, the interactions present the possibility of Twain hearing her own vocalizations. This may explain her maintaining interest in the interaction.

An alternate interpretation of her engagement, though, comes from examining the spectrograms of the original and broadcast exemplar; see Fig. S2 and SM D. Simply stated, the underwater loudspeaker broadcast playback was a very poor rendition of the animal’s powerful and throaty throp call—the actual exemplar. The broadcast exemplar was particularly lacking in low-frequency components and this might hint it was produced by a smaller and so younger humpback, whose vocal apparatus is necessarily smaller and so incapable of generating the low frequencies of an adult. If this was the case, then the extended interaction was not a time-delayed self-recognition study, but rather reflects the interest or empathy of a mother. (Twain had been sighted with a calf in July 2019 [14].)

## V. INTERACTIVE COMMUNICATION

Interactive playback is acknowledged as a powerful, if underused, exploratory tool for animal communication [46]. Not surprisingly, other fields also acknowledge and have established the benefits of interactive communication—benefits that bear technically on our results and hint at directions for future experimental protocols. For example, as developed in semiotic dynamics [47, Sec. V], interaction is key for a community to develop a collective communication system. Interaction also has practical benefits in greatly reducing the number of trials needed to monitor learning. On the mathematical side, computational learning theory shows that active learning— in which the student chooses queries that are potentially most informative—is faster and more parsimonious than passive learning—in which a teacher randomly selects and presents the facts to be learned independent of the state of the student [48]. Confronted with challenging field settings and logistics as with marine mammals, there is then the promise of experimental designs that afford such reductions in complexity and so could provide substantial savings in time and effort and improved accuracies.

The broad dichotomy between interaction and noninteraction bears directly on our acoustic study in two ways: First are the principled questions, How does one detect that two entities are interacting? Are they communicating? And, second is the question of experimental design for monitoring marine animals. Today, the predominant paradigm there relies on *passive acoustic monitoring* (PAM). Forced in large measure by the substantial difficulties of marine field work, one simply drops in from a vessel or anchors (e.g., to seafloor or animal [11]) the recording sensor (e.g., a hydrophone) to capture vocalizations, hopefully keyed to observed behaviors.

Recently, this has been augmented by using *playbacks*—broadcasting signals into the ocean via acoustic transducers; see SM A. That said, as an experimental protocol this is still passive one-way interaction as it broadcasts pre-recorded signals. This contrasts with what our experience leads us to advocate here: *interactive* playbacks in which a series of two-way interactions is sought where the component signaling is decided by each participant in real time. This has important consequences for experimental design as it greatly increases the demands on data acquisition, real-time audio processing, and the experimenters’ (at least) improvisational decision making.

Putting aside practical experimental difficulties, interactive playbacks pose substantial challenges when it comes to quantitatively establishing communicative exchanges from acoustic recordings. At first blush, three fields appear to offer appropriate methods for detecting the causality that is implied in interaction: mathematical statistics, multivariate information theory, and machine learning applied to the signals measured during interaction. Mathematical statistics frames the analysis as one of estimating joint and conditional distributions of the participants’ calls. Multivariate information theory [40] frames the analysis in terms of a bidirectional communication channel [41, 42, 49]—that is, two cross-coupled Shannon communication channels [50]. And, finally, machine learning frames this as validating one or another posited Bayesian network whose architecture captures the statistical dependence [51].

Our research methodology initially appealed to each of these to establish animal identity and human-animal information transmission. However, putting aside their requirements for very large data sets, we found that they each fail conceptually. Looking forward, the conclusion is that substantial theoretical innovations will be necessary for progress in analyzing interactive communication.

Stepping back, the results reported here strongly suggest moving beyond passive acoustic monitoring to playback studies. However, the extended interaction makes clear that this will not be sufficient. We propose *improvisational* interaction. In this, the field protocol requires a sound playback system that facilitates changes on the fly; much as a DJ would use to engage an audience. Also, of course, visual and sound information needs to be recorded. And, this leads one to consider a field-experiment setup equipped as a sound studio or video stage.

## VI. CONCLUSION

Let’s close by reviewing the results. The animal observed on 18 August animal was Twain: Fluke photo identifications, see. Fig. 4(a), identified by HappyWhale.com as Twain. The animal encountered on 19 August during interaction was Twain according to similar fluke identifications, see Fig. 4(b). The temporal coincidence of the visual and acoustic comparisons indicate that the subject animal was unique and was Twain.

The use of the 18 August throp call as a playback during the 19 August suggests the extended interactions could have been a kind of time-delayed vocal self-recognition study. Intentionally terminating the playbacks, the animal’s several repeated calls, and swimming away from the vessel bolster the interpretation of the animal’s intentional engagement. Thus, the experience, evidence, and analyses all point to the reported human-initiated playback-whale response being a simplified form of interactive communication. While, the interaction and our quantitative analyses leave a number of questions open, they do offer a hopeful, but daunting challenge to future interactive studies of cetacean communication.

After reviewing the long history of cetacean playback studies—see SM A—to the best of our knowledge the extended series of acoustic interactions with Twain is unique. While a hopeful occurrence, it poses more questions than answers—Was it a conversation between human operators (with their improvisationally-determined timings of pre-recorded playbacks) and Twain the humpback? What quantitative criteria can be established to demonstrate communicative interactions in terms of conditional statistical dependence, information flow and exchange, and intentional, causal interaction?

One practical consequence for future field research suggested by the results is that interactive playbacks require technical support for improvisation. Genuine communication resides in improvisation. More directly, one cannot hope to understand animal communication, in whatever form, without experimental protocols that support sonic improvisation and without a firm theoretical basis for quantitatively analyzing the resulting information flow, contingency, and causality.

Success in answering these and related questions will bear directly on future explorations of interspecies communication and, more generally, human appreciation of nonhuman intelligence. While analogous interactions are familiar in other species, the abiding question in designing future studies is how to reliably establish such communicative interactions with cetaceans in natural, noncaptive environments.

In the last three hours of the day, God sits and plays with the Leviathan, as is written: “you made the Leviathan in order to play with it”. *Talmud, Avodah Zarah*

## ADDITIONAL INFORMATION

Supplementary Materials are available for this article.

Correspondence and requests for materials should be addressed to the first author.

## DATA AND MATERIALS AVAILABILITY

All data needed to evaluate the conclusions in the paper are available from the first author upon reasonable request.

## ACKNOWLEDGMENTS

All investigations conducted under National Marine Fisheries Service Research Permit #19703. The authors are deeply indebted to Don and Denise Bermant (M/Y Blue Pearl, Vancouver, British Columbia) and Marc Choquette (Captain) and Collette Costa (M/V Glacier Seal, Juneau, Alaska) for their generosity and expert seamanship. They are also grateful to Tony Gilbert of SeaKeepers.org and Laurance Doyle and Debbie Kolyer of the SETI Institute. Jodi Frediani, Aza Raskin, Brit Selvitelle, Clark Snodgrass, and Katie Zacarian provided observational support. We thank them all and Nicolas Brodu, David Dunn, Kelly Finn, Yolande Harris, Ari Friedlaender, Josie Hubbard, Brenda McCowan, Fred Sharpe, and Andy Szabo for helpful conversations. Finally, we thank the Alaska Whale Foundation for its kind hospitality during visits to Warm Springs, Baranof Island.

## Field Contributions

J.P.C., Brenda McCowan (B.M.C.), and Fred Sharpe (F.S.) conceived the experiment. J.P.C. and F.S. designed and assembled the experimental apparatus; J.P.C. wrote the control/measurement software; J.P.C. and B.M.C. performed the acoustic playbacks and hydrophone recordings. F.S. directed the top-deck observation team, including Josie Hubbard (J.A.H.) who recorded the camcorder video. J.P.C., B.M.C., and F.S. provided the recording and playback equipment.

## Author contributions

J.P.C. and A.M.J. analyzed the experimental data; J.P.C. composed the videos and audio, including hydrophone recordings; J.P.C. and A.M.J. performed statistical analysis and theoretical modeling; J.P.C. and A.M.J. performed numerical simulations. J.P.C. and A.M.J. prepared the manuscript and discussed the results and their implications.

## Supplementary Materials

The Supplementary Materials recount the relevant natural history of humpback whales, recent history of acoustic interaction studies, experiment design and protocols for interactive playbacks, 18 August 2022 sighting of humpback whale Twain, details of the 19 August 2022 acoustic interactions with her, determining Twain’s identity from hydrophone recordings and photographic sighting, background on applying the Wasserstein metric to power spectral densities, playback and response timings during the encounter, statistical contingency of those interactions, and pointers to the playback-response audio-video recordings.

## Appendix A: Humpback Whale Natural History and Behavioral Response Studies

The main text provided a brief natural history of humpback whales and a brief history of acoustic playback studies. The following provides further details, many of which are important to a fuller understanding and interpretation of the extended interactions with Twain.

Modern cetacea (whales, dolphins, and porpoises) descended from land-dwelling, carnivorous ungulates that entered the shallows of the warm Tethys Sea in the Eocene (57*−* 35 million years ago) and never returned to land [52–54]. Since then two orders of whale evolved: the odontoceti (toothed whales) and the mysticeti (baleen whales). The mysticeti evolved baleen around 25 million years ago, losing their teeth.Today, the mysticete species of humpback whales (*Megaptera novaeangliae*) are found in all the planet’s oceans. They migrate annually between high-latitude coastal summer feeding grounds and low-latitude breeding and calving grounds in shallow waters. Of the North Pacific population, a significant fraction winters each year in waters surrounding the major Hawaiian islands and returns to Alaskan waters during the summer months to feed [17–19]. The social behavior of humpback whales while in Hawaiian waters is largely related to reproduction.

Humpbacks are one of the most vocal whale species. They generate sounds underwater by passing air between the lungs and the laryngeal sack through vestigial vocal cords.

Whales have a special auditory view of their environment due to their successfully coupling an air-adapted mammalian ear to underwater sound [55]. First, they have the only mammalian ears fully adapted to underwater hearing. Second, they employ the broadest acoustic range of any known mammal group. Considering that cetaceans function wholly in water, a dense medium in which light attenuates far faster than sound, it is not surprising that sound is believed to be whale’s primary medium for supporting sensory and communication channels.

To hear a sound equally well in water and in air an intensity-based mammal ear requires sound pressures in water sixty times that in air. Baleen vocalizations fall in the sonic to infrasonic range (12 Hz-3 kHz) [56].

Most important, both odontocete and mysticete auditory innervation densities are significantly greater than those of other mammals. Baleen whales, in particular, can extract complex data from low to infrasonic signals; i.e., they may be infrasonic echolocators. Cetaceans have a 10-11 octave functional hearing range compared to human’s 8-9 octave range with a maximal functional high frequency capacity of 20 to 30 kHz. The consensus is that mysticete ears are adapted for good low to infrasonic hearing.

Most cetaceans are social animals [57]. Humpback social behavior appears to be substantially moderated by vocalizations of two kinds. The first are social calls: short duration (2-20s), occurring in sudden flurries of sounds interspersed by long silences, and produced in groups of 3 or more adults [24, 58]. The second are songs: long (minutes to hours), extended complex patterns, and rhythmic and continuous [59, 60]. The long, complexly structured songs are heard throughout the winter breeding grounds for the duration of the season. The singer is normally a lone male whale [19, 61, 62]. Under the right conditions songs can be heard for hundreds of kilometers.

Social sounds, in contrast, lack the complex structure of song and, unlike song, have been heard only in the context of large, surface-active pods containing multiple escorts [24, 58]. Humpbacks appear to be less vocal in their summer feeding grounds than in the winter grounds [63].

Humpback whales employ a unique and complex foraging behavior—bubble-net feeding—that involves expelling air underwater through their blowhole to form a vertical cylinder of bubbles around prey to contain and possibly compact them. Often the construction of the bubble-net is coordinated in groups and feeding involves changing rolls from one event to the next [64].

Given their water environment and behaviors that keep them below the surface for extended periods of time, acoustic monitoring of humpbacks using hydrophones is a primary method of investigation [21]. In addition to passive monitoring by hydrophone there is a long history of sound playback studies going back to the 1950s to probe whale behavior responses to sound. Playback experiments were first used with cetaceans in 1952 to determine the upper limits of hearing of the bottlenose porpoise *Tursiops truncatus* [65, 66]. Playback of conspecific animal sounds has been used since 1961 as a means of demonstrating vocal exchanges between dolphins. For example, sonic stimuli from one T. truncatus immediately elicited whistles and click trains from another, isolated animal of the same species [67]. Moreover, certain “signature” whistles were specific to the individual producing them.

Playback studies showed that whales respond to recordings of their own sounds in complex, context-dependent ways. Both syntax (combining individual sounds into a series) and context were important in conveying information by playbacks [23]. For example, southern right whales *Eubalaena australis* approached an underwater loudspeaker broadcasting right whale sounds or human imitations of right whale sounds. They increased their own rate of vocalization during such playbacks. However, they swam away and made relatively few sounds in response to playbacks of water noise, 200 Hz tones, and humpback whale sounds. Thus, southern right whales can differentiate between conspecific sounds and other sounds [68].

Yet another type of experiment used the playback of sounds of one cetacean species to another. For example, gray whales immediately took escape behaviors to avoid the source of playbacks of orcas, a known predator [69].

More germane to the present report, the differential response of humpbacks to social sounds and to song was studied during the breeding and calving season off the island of Maul, Hawai’i [24]. Singing whales stopped singing when exposed to playback of either songs recorded from lone whales or social sounds recorded from humpback whale groups in which males were fighting. Cows with calves and groups of three of more adults exposed to social sounds moved away. Whales exposed to playback of song moved away.

Most dramatically, adult humpbacks exposed to social sounds approached the playback vessel: it is “impressive to an observer in a 5 m boat when a 15 m whale charges the boat” [24]. The whales that charged swam at the surface approaching to less than 5 m from the playback speaker and then “swam around the boat in ever widening circles, occasionally making another pass under it. Although charging whales came as close as 2 m to the speaker, they never touched it”. A later study encountered similar response [70]:

The most striking behavioral response to sound playback was the rapid approach of one or more individual whales to the playback vessel. The approaching animals typically dove close to the vessel, then either swam rapidly past the boat to one side or directly under it. Occasionally, they could be seen passing under the speaker. At other times, they surfaced close by and circled the vessel.

Attraction under these playback paradigms might be traceable to the contextual novelty of the sound—one not normally heard during the winter season—and that novelty induces approach. Hearing a feeding call might indicate recognizing the latter as not only contextually novel, but coming from a conspecific.

Continued studies of Hawaiian humpback whales on their winter calving grounds revealed that social sounds occurred almost exclusively in whale groups and were rarely heard near single whales, pairs, or cow-calf groups. Moreover, social sounds were heard rarely in nonaggressive situations and vocalization increased markedly with the appearance of new whales. Altogether, these observations were interpreted as providing evidence for the social role of humpback vocalizations [58].

Reference [71] carried out behavioral response studies with humpback whales to test the response of groups to one recording of conspecific social sounds and an artificially generated tone stimulus. The whales responded differently to the two stimuli. The response to social sounds was highly variable and depended on social group composition.

When designing such studies, an initial challenge is to select which of the 30 or so humpback social calls to broadcast [12, 26, 72]. A natural guess is that the most frequently produced calls in an animal’s repertoire (throps and grunts) are important. And so, they make suitable initial candidates to elicit responses and explore function and meaning. The acoustic interactions we report used the humpback throp call (sometimes described as a “whup” call). Purportedly, this is a social greeting call [12]. The vocalization starts out as a low-frequency amplitude-modulated call (“growl”) that ends with an increasing-frequency sweep. Humpback whales vocalize throps across all oceans. The call has persisted across generations. The calls are believed to be innate in this species [25–29]. The current consensus is that their communicative role is that of contact or greeting.

**FIG. S1:**
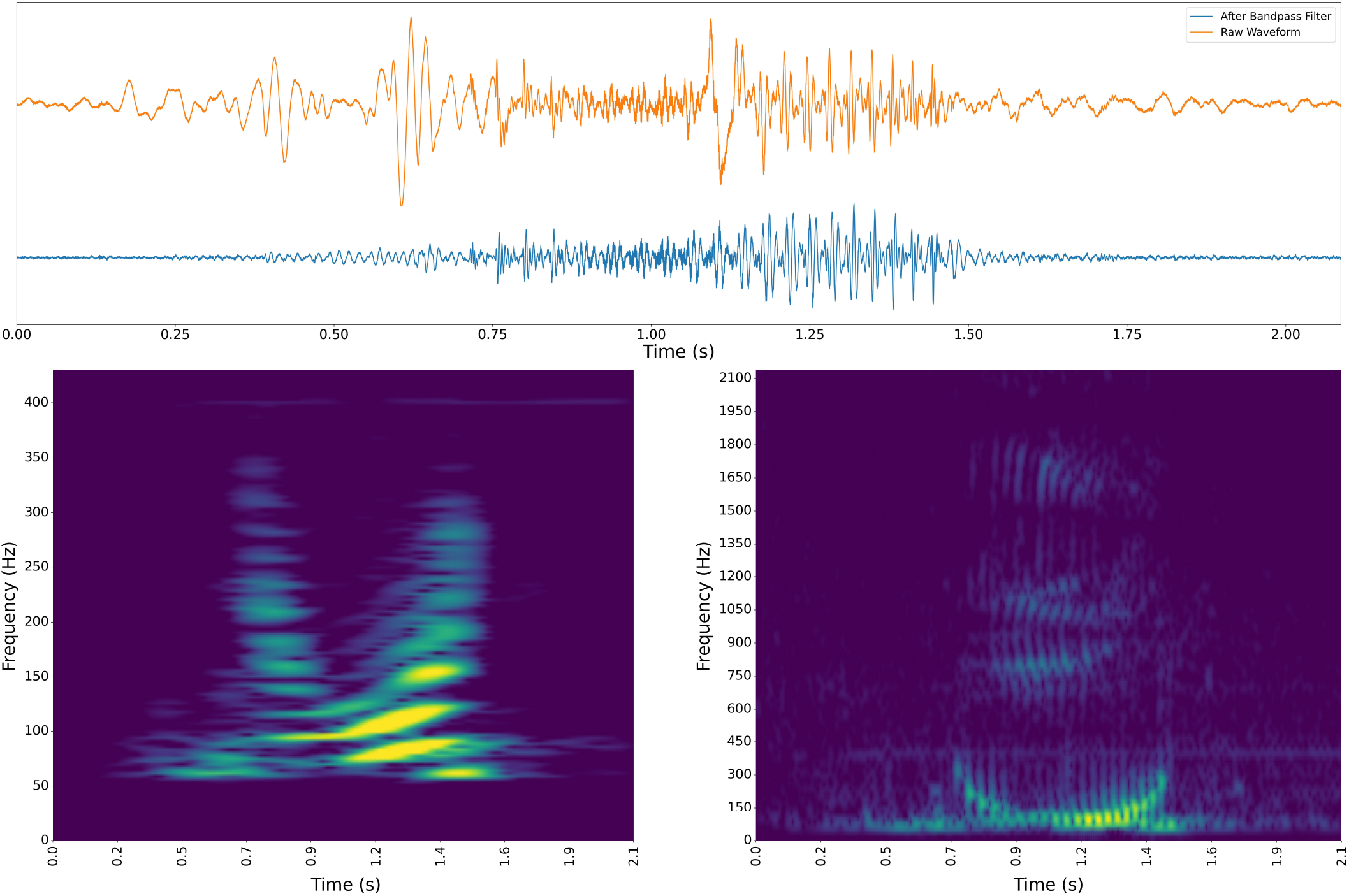
Throp Exemplar: (Top) Waveform before (above, orange) and after (below, blue) bandpass filtering. Waveform was filtered with a 20th-order digital Chebyshev type II filter with critical frequencies 40 Hz and 5000 Hz. (Bottom) Spectrograms at low frequencies (left) and at high frequencies (right). The different texturing in the two spectrograms indicates the use of two different window widths in the short-time Fourier transform (STFT). The window width of an STFT determines the resolution of the spectrogram. When the window is wide, as on the left (*∼*40 ms), the STFT gives good frequency resolution and we may see the harmonics of the whale’s throp. When the window is narrow, as on the right (*∼*4ms), the STFT instead has sharp time resolution, showing the trilling beats of the throp sound, but blurs out the frequencies present. The frequency-time uncertainty principle of STFTs is a familiar phenomenon; noted even in the earliest speech sound spectrography analyses [31, 32].

The long history of playback studies contrasts with the present results. Specifically, interactions with Twain are atypical in light of previous reports of humpback social vocalizations. First, previously social sounds occurred largely in groups containing three or more whales. Social sounds were infrequently heard near single whales, pairs, or cow-calf groups. Our playback interactions involved only one vocalizing animal—Twain. Others were present, but did not vocalize. Second, the interaction occurred in humpback summer feeding grounds—southeast Alaska, not as previously in their Hawai’i winter calving grounds. Third, in the summer feeding grounds little to no song is vocalized and little aggression is observed, likely due to the absence there of reproduction activity and mating competition. Fourth, previously social sounds were heard rarely in nonaggressive situations. In contrast, the Twain throp interactions took place in a nonaggressive setting. Fifth, previously interactions were reported to fall into two classes of (i) exclusively rapid approach to the broadcast vessel or (ii) nonapproach (move away from vessel). Twain stayed near the vessel for a half hour. Finally, previous studies did not employ mutual acoustic exchanges. Certainly, in previous studies, what communication there was was not interactive, in contrast to the repeated turn-taking interactions with Twain.

## Appendix B: Experiment Design and Protocol for Interactive Communication

To stimulate sonic interactions during the playback trials a Lubell System 3400 was deployed, including an underwater loudspeaker (Lubell LL916C-025’) driven by a 60W audio mixer/amplifier (Lubell modified TOA CA160 amplifier) powered by a 12V marine battery. From the vessel’s main deck, a laptop computer (Apple MacBook Pro 15”) drove the amplifier with a collection of prerecorded natural and synthetic sounds.

To capture both the playbacks and animal responses a single omnidirectional hydrophone was deployed—a Model 54, Cetacean Research Technology (Seattle, WA) with a frequency response of 0.02*−* 50 kHz and effective sensitivity of *−*169 dB re 1v/uPa). The hydrophone signal was recorded by a separate laptop (Macintosh MacBook Pro 15”) also on the main deck. Playback and recording sample rates were set at 44.1 kHz with 16 bits per sample. See Fig. 1 for the overall setup. The computer operators wore stereo headphones (Sony MDR-6) to mask environmental noise, concentrate on the hydrophone signals, and be sensitive to distant and spatially-distributed animal vocalizations. Vessel location and orientation was monitored via GPS (Bad Elf 2200 GPS Pro).

The speaker and hydrophone were independently suspended at a depth of approximately 20 feet below the vessel roughly 10*−* 20 feet apart. That noted, strong local ocean currents and wind-forced vessel drift varied this separation considerably on occasion.During playback experiment trials two operators—J. Crutchfield (JPC) and B. McCowan (BMC)—were inside on the main deck at the electronic equipment station running the playbacks (using Raven v. 1.6) and the recordings (via Audacity v. 3.0.2). Observers—J. Hubbard (JAH), F. Sharpe (FS), and C. Snodgrass (CSS)—were stationed on the upper deck with camcorders (video and voice recording observations), various still cameras, laser range finders (TideWe HR-F1000), binoculars (NIKON 7 × 50 7.2 deg CF WP COMPASS), and notebooks to write out narratives of visual sightings.

On most daily excursions, upon sighting nearby humpbacks and approaching them (*≈* 100m), for each experimental trial a simple three-part protocol was implemented: recording for 10 minutes as a control *before* period, then broadcasting playbacks of various kinds for 10 minutes—the *during* period—and, finally, continued recording for another *after* control period without playbacks. Repeatedly, these attempts were rebuffed by nearby whales who met them with utter silence. Given active vocalizations by the southeast Alaska humpback population, noted in prior reports and captured by us during the voyage, the complete lack of acoustic interaction was notable. This will be reported on elsewhere.

Despite the agreed-upon experimental protocol, the interaction with Twain was so markedly surprising, on its own and in contrast with prior noninteractions, that JPC deviated from the protocol as soon as the (as-yet unknown) animal closely approached the vessel. Specifically, the “during” playback broadcasts substantially exceeded the agreed-upon protocol—20 minutes rather than 10. We now recount additional details of the encounter beyond those provided in the main text.

## Appendix C: Twain Identity and Throp Call (18 August 2021)

Twain was sighted on 18 August 2021 (14:38:40 PST) during planned excursions around Frederick Sound. Several photographs captured her fluke prior to diving. See Fig. 4(a). It was only later on the following day that this photograph was matched on HappyWhale.com to a humpback whale Twain sighted many years prior; see, for example, Fig. 4(c).

Also, on 18 August, a number of hydrophone recordings and several interaction experiment trials were performed. On reviewing these recordings a particularly clear call was identified, known as a *throp* social call [63, 73]. That recording was minimally contaminated with environment noise (low sea-state, minimal vessel noise) and other animal vocalizations. The call employed as a playback was a medium-frequency, frequency-modulated pulsed call. We refer to this as the *throp exemplar* and denote it *E*0 elsewhere. Figure S1 gives its waveform (raw and bandpass filtered) and spectrogram.

## Appendix D: Twain Interactive Acoustic Encounter (19 August 2021)

A clear and distinct response to a playback broadcast occurred on the first experiment trial of 19 August, soon after the R/V Glacier Seal departed Hobart Bay anchorage mid-morning. The winds were calm and seas were smooth with moderate-to-heavy fog with visibility less than a quarter mile. As the vessel cruised south seeking experiment opportunities in east Frederick Sound, the first whale sighting occurred northeast of and near Round Rock. The vessel stopped and was quieted.

**FIG. S2:**
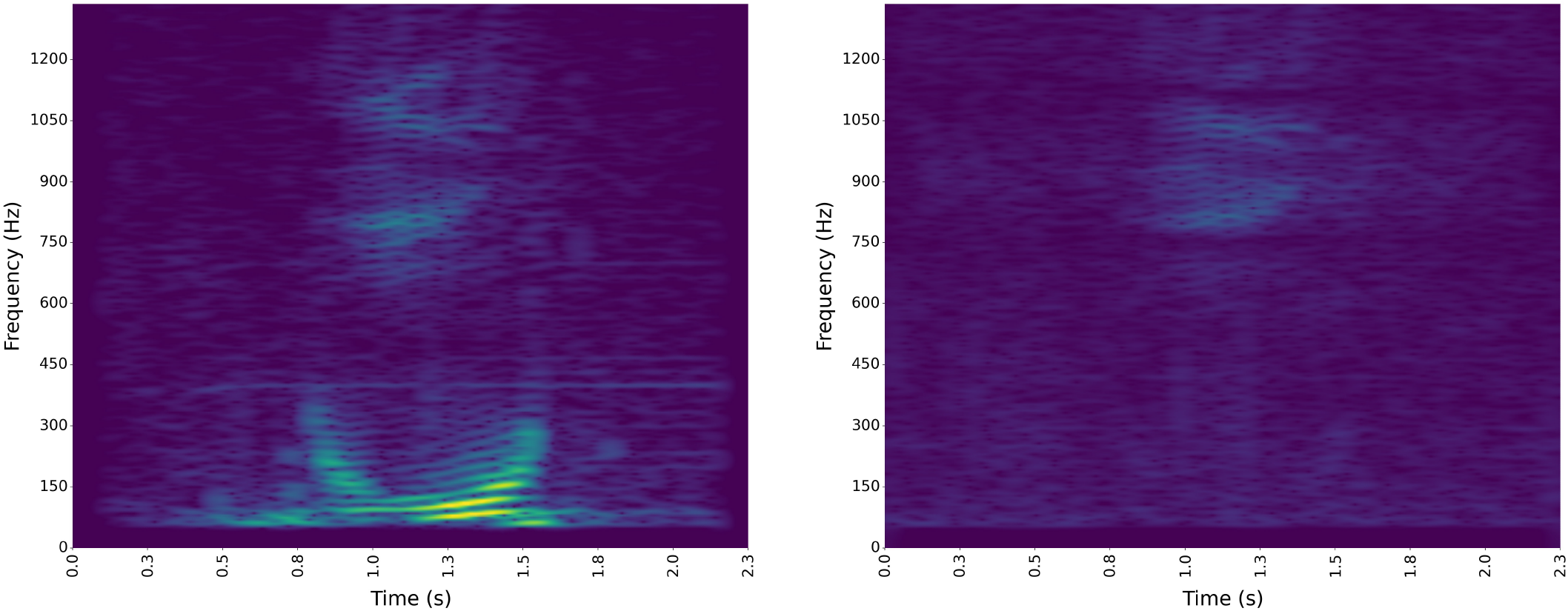
(Left) Spectrogram of throp call *exemplar* recorded on 18 August. (Right) Spectrogram of throp *playback*— hydrophone recording of the exemplar as broadcast 19 August. The integrated intensity in each is a proxy for the total acoustic energy in each. The animal’s throp (Left) is substantially more powerful (easily heard as much louder) than the underwater speaker playback (Right) that lacks much power at low frequencies. The stark difference raises concerns about what Twain actually heard during the extended interactions.

Figure S2 (Left) gives the throp exemplar spectrogram to compare against the recording of its underwater loudspeaker playback on the 19^th^, Fig. S2 (Right). The differences are notable with the exemplar broadcast being markedly less powerful and lacking substantial low frequency components, due to the spectral response of the underwater loudspeaker. Nonetheless, Twain’s own throp call was played back to her on the 19^th^.

Timings of the human-initiated playbacks were determined on-the-fly by the operators (JPC and BMC), allowing for improvisational control to probe for novel responses, turn-taking, and temporal mirroring. The videographer (JAH) activated a video camcorder (Sony Handicam NEX-VG10) at the beginning of the interaction and synced with the Audacity-controlled recording by recording its window on the laptop screen. Topside, the videographer and observers (FS and CSS) were blind to playback treatment. However, complete sound isolation was compromised by the low environmental noise and the underwater speaker’s close placement.

During the before-period, visibility improved to several miles. The first activity observed was prey and surface bubbles. This was followed by vigorous surface lunge near prey at 100 m. This included a jaw slap that later facilitated synchronizing the hydrophone recording and topside observations. The whale then continued slowly traveling in the vessel’s direction. The first exemplar playback *P*_0_ was broadcast to an individual estimated at 40 m. A second playback *P*_1_ was broadcast. The animal then rose to take its first breath, then sinking back below the surface. After the third throp call *P*_2_ was broadcast the animal gave its first response *R*_0_.

Figure 5 (Left) presents the spectrogram of the throp exemplar *E*_0_ and Fig. 5 (Right) that of several of Twain’s responses—*R*_0_*−, R*_19_, and *R*_35_—during interactions. Figure S3 (Above) presents Twain’s response throps *R*_0_ − *R*_35_ lined up to show their close uniformity, once the overall power in each response is accounted for. Twain’s responses are relatively uniform, especially in duration, pulse rate, and number of pulses. However, the initial pulse in the response is variable in timing and power. There is also variation in the harmonics. However, repeatedly using a single playback exemplar provided no signal novelty that might have stimulated Twain, recall Fig. 6 (Below).Figure S4 presents the sequence of the playback (*P*_*i*_) and response (*R*_*j*_) energy spectra across the extended interaction. The playback spectra are plotted in red; responses in blue. *E*_0_ there labels the exemplar spectrogram. The similarity between the exemplar and responses is relatively clear. What is notable, as indicated elsewhere with other methods, is the difference between the playbacks—the hydrophone recording of the broadcast exemplar—and the responses and the exemplar itself. To even see the playback energy spectra in the plots required substantial amplification. That is, if the playbacks were normalized in the same manner as the responses, the playback spectra details would be barely visible.

These observations point to the sonic paucity of the broadcast playbacks. And, this low quality is due to inadequacies of the underwater loudspeaker and audio amplifier. Louder playbacks led to audible distortion in the playbacks. This offers up a number of questions. Not the least of which is, what was Twain hearing in the playbacks?

**FIG. S3:**
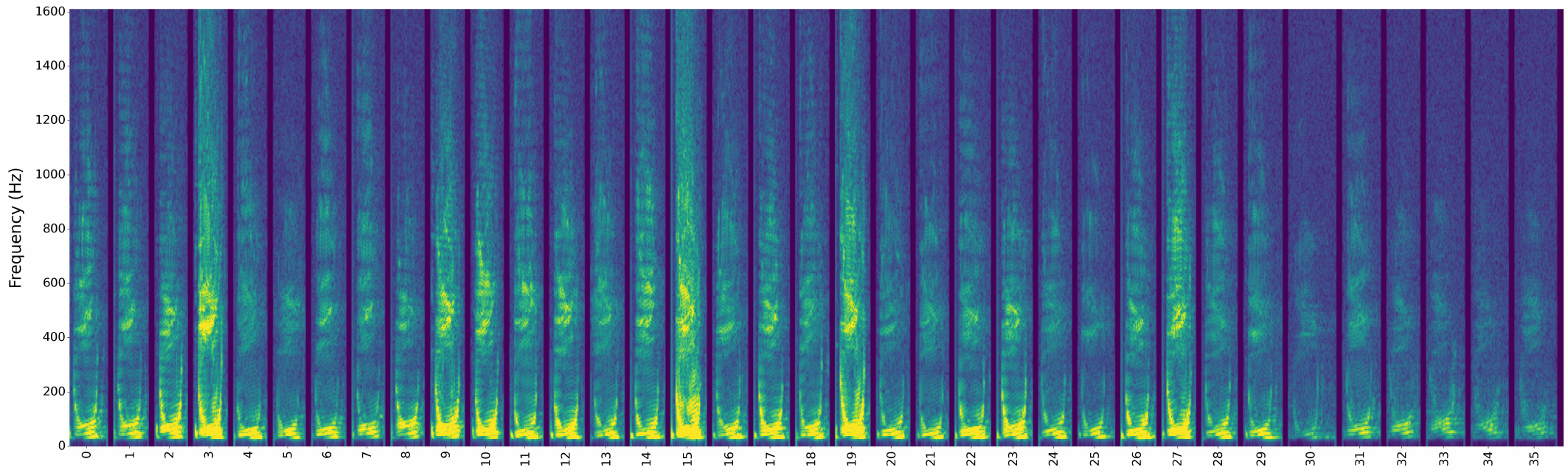
Acoustic identity across interactions: Spectrograms of Twain’s throp responses *R*_0_ − *R*_35_.

Why did she respond, given the seemingly large spectral differences? Noting the bulk of the playback spectra is at relatively high frequencies, perhaps the throp playbacks seemed to be those from a juvenile humpback—juveniles being smaller and so having smaller vocal generating organs.

Finally, looking across the responses, it is clear that, while similar, they are also spectrally distinct at some level. In the differences, is there semantic information that Twain used to try to communicate something novel? Or, perhaps the differences simply reflect variable undersea conditions. At this point, we have been unable to address these questions using available signal analysis, information-theoretic, and machine learning tools.

### 1. Playback and Response Timings

Table S1 gives the measured timings between playback and animal response from the hydrophone recording in Fig. 2. The times quoted are the beginning of each sound—playback *P*_*i*_ or response *R*_*i*_. The time ∆ between sonic events is given, along with various annotations as to surface observations.

### 2. Conversational Contingency

The extended interaction was a series of sonic events—almost exclusively turn-taking vocalizations between human-initiated playbacks *P*_*i*_ met with animal responses *R*_*j*_. Thus, the hydrophone recording is a time series of two kinds of event separated by time intervals. One question this observation brings up is whether there is communicative information or perhaps contingency in the event series.

A standard method method for analyzing such time series was introduced by the meteorologist Ed Lorenz in 1963 to address this question in the temporal evolution of fluid turbulence [74]. He studied, in particular, a three-mode simplification of the hydrodynamic equations—a set of 3 first-order differential equations—showing they generated what is now called deterministic chaos. That is, though apparently noisy, there was deterministic dependency between future observations and past observations.

The question Lorenz posed was: Is the next event a function of previous events? If we denote an event at time *t* by *x*_*t*_, this question becomes one of determining whether or not there is a function *f* (·) such that:

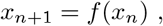

or, if there might be longer dependencies:

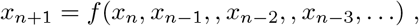

where the values are continuous: *xn ∈* R. If there is not a deterministic function *f* (·), then one allows for noise, using:

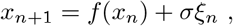

**TABLE S1:** Interaction timings of playbacks (*P*_*i*_) and animal responses (*R*_*j*_). Times here relative to first playback *P*_0_in Fig. 2.

| Time | $\Delta$ | Sound | Comment | Time | $\Delta$ | Sound | Comment |
| --- | --- | --- | --- | --- | --- | --- | --- |
| | | | | 08:20 | 00:16 | $P_{20}$ | |
| 00:00 | | $P_0$ | First Playback | 08:24 | 00:03 | $R_{18}$ | |
| 00:18 | 00:18 | $P_1$ | | 08:32 | 00:08 | $P_{21}$ | |
| 00:44 | 00:25 | $P_2$ | | 08:53 | 00:21 | $R_{19}$ | |
| 01:21 | 00:37 | $R_0$ | First Response | 09:07 | 00:14 | $P_{22}$ | |
| 01:32 | 00:10 | $P_3$ | | 09:16 | 00:08 | $R_{20}$ | |
| 01:39 | 00:06 | $R_1$ | | 09:32 | 00:15 | $R_{21}$ | Surface |
| 01:47 | 00:07 | $P_4$ | | | | | Dive |
| 01:59 | 00:12 | $R_2$ | | 09:39 | 00:06 | $P_{23}$ | |
| 02:09 | 00:10 | $P_5$ | | 10:07 | 00:27 | $P_{24}$ | |
| 02:18 | 00:08 | $R_3$ | | 10:19 | 00:12 | $R_{22}$ | |
| 02:27 | 00:09 | $P_6$ | | 10:28 | 00:09 | $P_{25}$ | |
| 02:45 | 00:17 | $R_4$ | | 10:40 | 00:11 | $R_{23}$ | |
| 02:56 | 00:11 | $R_5$ | | 10:50 | 00:09 | $P_{26}$ | |
| 03:04 | 00:08 | $P_7$ | | 11:22 | 00:32 | $R_{24}$ | |
| 03:16 | 00:11 | $R_6$ | | 11:40 | 00:17 | $P_{27}$ | |
| 03:27 | 00:11 | $P_8$ | | 11:44 | 00:03 | $R_{25}$ | |
| 03:38 | 00:10 | $R_7$ | | | | | Surface |
| 03:48 | 00:09 | $P_9$ | | 11:49 | 00:04 | $P_{28}$ | |
| 04:05 | 00:17 | $R_8$ | | 12:02 | | | Breathing |
| 04:15 | 00:09 | $P_{10}$ | | 12:29 | 00:39 | $R_{26}$ | |
| 04:29 | 00:13 | $R_9$ | | 12:49 | 00:20 | $P_{29}$ | |
| 04:40 | 00:11 | $P_{11}$ | | 12:52 | 00:03 | $R_{27}$ | |
| 04:52 | 00:11 | $R_{10}$ | | 13:03 | 00:10 | $P_{30}$ | |
| 05:02 | 00:09 | $P_{12}$ | | 13:30 | 00:27 | $R_{28}$ | |
| 05:12 | 00:10 | $R_{11}$ | | 13:41 | 00:10 | $P_{31}$ | |
| 05:21 | 00:09 | $P_{13}$ | | 13:56 | 00:15 | $R_{29}$ | |
| 05:26 | 00:04 | $R_{12}$ | | | | | Surface, Move away |
| 05:32 | 00:06 | $P_{14}$ | | 13:58 | 00:02 | $P_{32}$ | |
| 05:55 | 00:23 | $R_{13}$ | | 14:25 | 00:27 | $P_{33}$ | Surface, Move away |
| 06:06 | 00:11 | $P_{15}$ | | 14:50 | 00:24 | $R_{30}$ | |
| 06:21 | 00:14 | $R_{14}$ | | 15:02 | 00:12 | $P_{34}$ | |
| 06:34 | 00:12 | $P_{16}$ | | | | | Surface, Move away |
| 06:51 | | | Breathing | 15:32 | 00:29 | $R_{31}$ | |
| 07:09 | 00:35 | $P_{17}$ | | 15:43 | 00:10 | $P_{35}$ | Last Playback |
| 07:20 | 00:11 | $R_{15}$ | | 16:00 | 00:17 | $R_{32}$ | |
| 07:32 | 00:11 | $P_{18}$ | | 16:40 | 00:40 | $R_{33}$ | |
| 07:37 | 00:05 | $R_{16}$ | | 17:21 | 00:40 | $R_{34}$ | |
| 07:46 | 00:08 | $P_{19}$ | | 18:02 | 00:41 | $R_{35}$ | Last Response |
| 08:04 | 00:18 | $R_{17}$ | | | | | |

where *ξn* is a unit-standard-deviation Gaussian random variable and *σ* a parameter that sets the *noise level*. In time series analysis and dynamical systems theory this is the data analysis method of *return maps* used to search for contingencies in discrete-time series of continuous-valued events.

Notably here, dynamical-systems return-map analyses [74, 75] have already been used to good effect in marine biology to discover rhythmic modulation of sperm whale echolocation click-train vocalizations [76]. There, naturally enough, the return-map events consisted of clicks and so the resulting time series was of “interclick intervals”. Here, instead we used “intercall intervals”.

More to the point, in the extended interaction there are two kinds of events, where the kinds form a set *V* = *{P, R} P* denoting a playback vocalization and *R* a response vocalization. That is, vocalization *v ∈ V*. In addition, the *i*^th^ event *e*_*i*_ has a nonnegative time of occurrence *t*_*i*_ ∈ ℝ_+_. So, in fact, events *e* here are pairs *e*_*i*_ = (*v*_*i*_, *t*_*i*_) and the event time series is **e** = (*e*_0_, *e*_1_, *e*_2_, …, *e*_*M*_). In this, *M* is the length of the event time series.

**FIG. S4:**
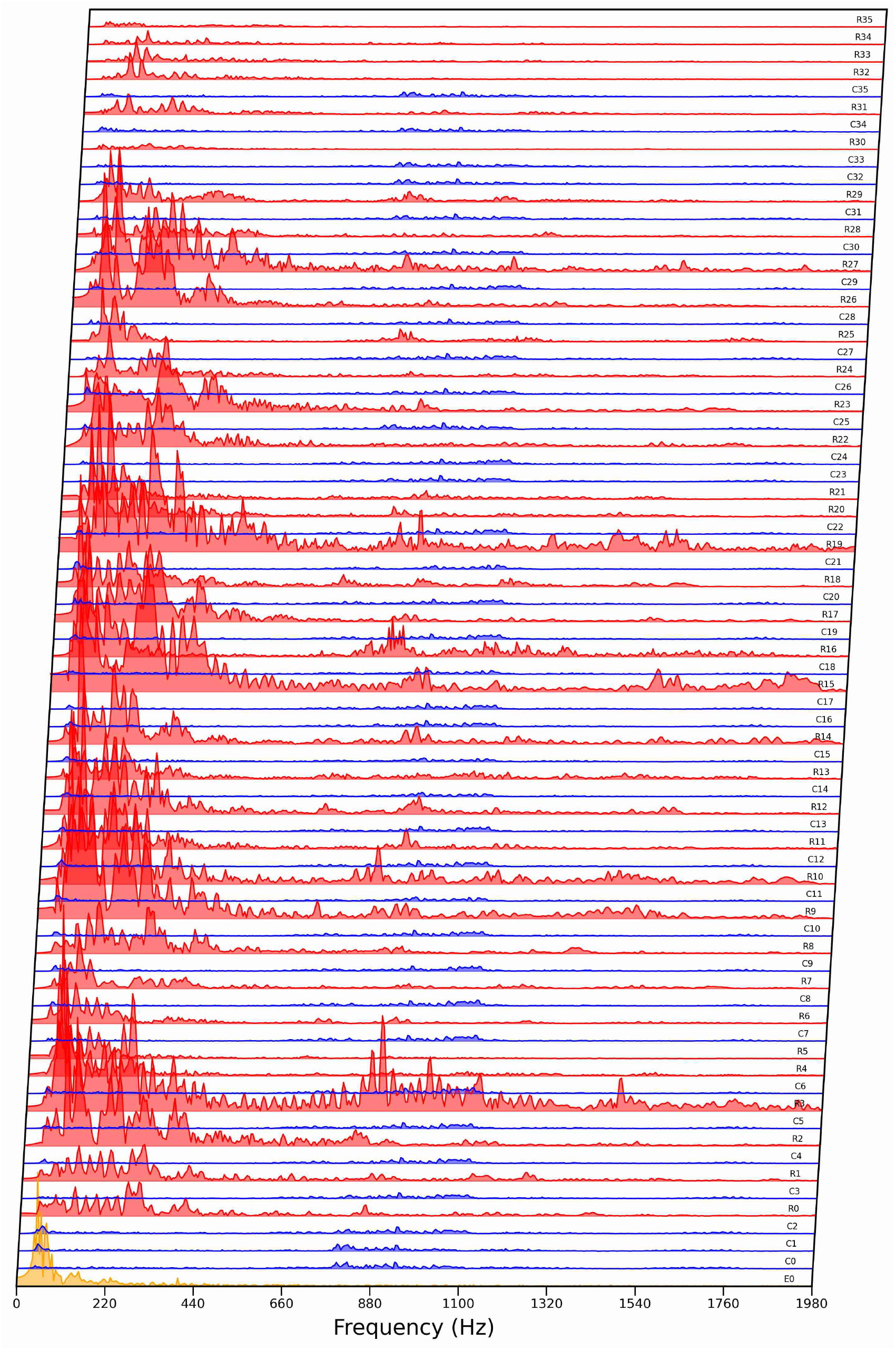
Time sequence of playback (blue) and response (red) spectral transforms from interaction beginning (bottom) with three playbacks (P0-P2) before the animal first responds (R0) to its end (bottom) with the last playback (P35) and Twain’s final responses (R33-R35). The exemplar, shown in yellow, has a spectral profile more similar to the responses.

**FIG. S5:**
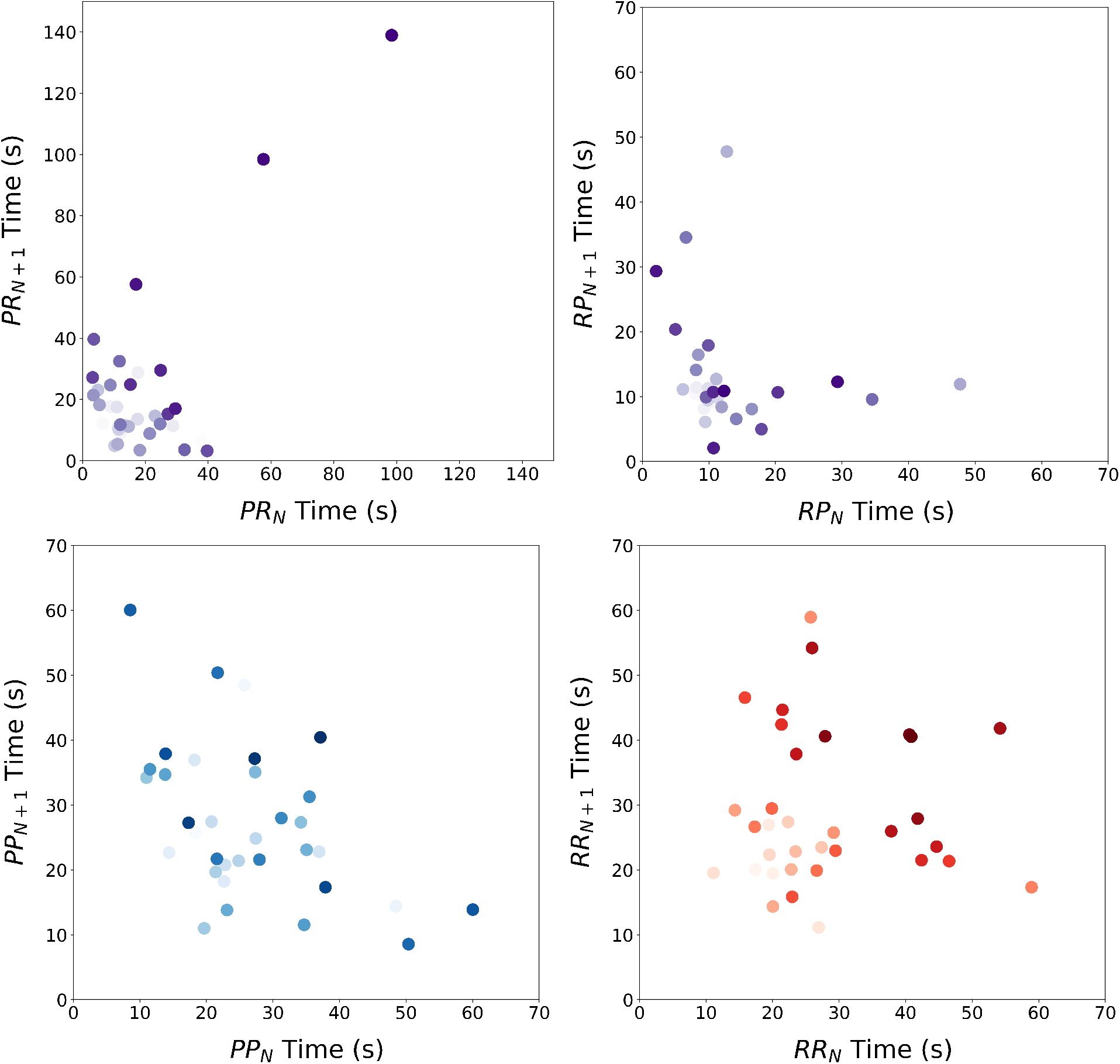
Interaction event return maps: (Top Left) Playback-to-response (PR), (Top Right) Response-to-playback (RP), (Bottom Left) Playback-to-playback (PP), and (Bottom Right) Response-to-response (RR). Dot color indexes the time of occurrence with lighter being early, darker late.

Stochastic process theory refers to such discrete-value time-of-occurrence data—a discrete sequence of continuous-valued events—as realizations of a *semi-Markov process*. Reference [77] shows what kinds of generative processes these are in terms of their minimal generating mechanism and Ref. [78] introduces an algorithm for inferring the latter from data.

The encounter consists of only *M* = 72 events. Due to this small sample size, here we only probe the existence of functional contingency by plotting event return maps to visually capture *f* (·). Figure S5 (Top) explores temporal dependencies in the playback-to-response (PR) and response-to-playback (RP) event time series that capture the interaction dependencies. Figure S5 (Bottom) explores temporal dependencies in the playback-to-playback (PP) and response-to-response (RR) event time series to capture possible long-time turn-taking dependencies.·

We see that there is little evidence of deterministic dependency. The effective return maps *f* (·) are quite random. There is, however, weak statistical contingency that appears as a lack of data points in both (i) the upper right area of long interevent times and (ii) the lower left areas of short interevent times. Apparently, both the operators and the animal (i) did not respond too quickly and (ii) typically responded within a minute to each other.The exception to the generally nondeterministic dependency shows up in the *PR* times: As Twain moves away, having heard the last broadcast playback, the events are only responses and the time to last playback grows with each of Twain’s uninvited responses. It would seem that when the human-initiated playbacks ended, her expected time to hear a playback went over a minute. From this she correctly decided the interaction was over and swam away, though calling several more times. Were those attempts to solicit further human playbacks? That aside, apparently the three further tries at interaction were enough and she stopped vocalizing.

**FIG. S6:**
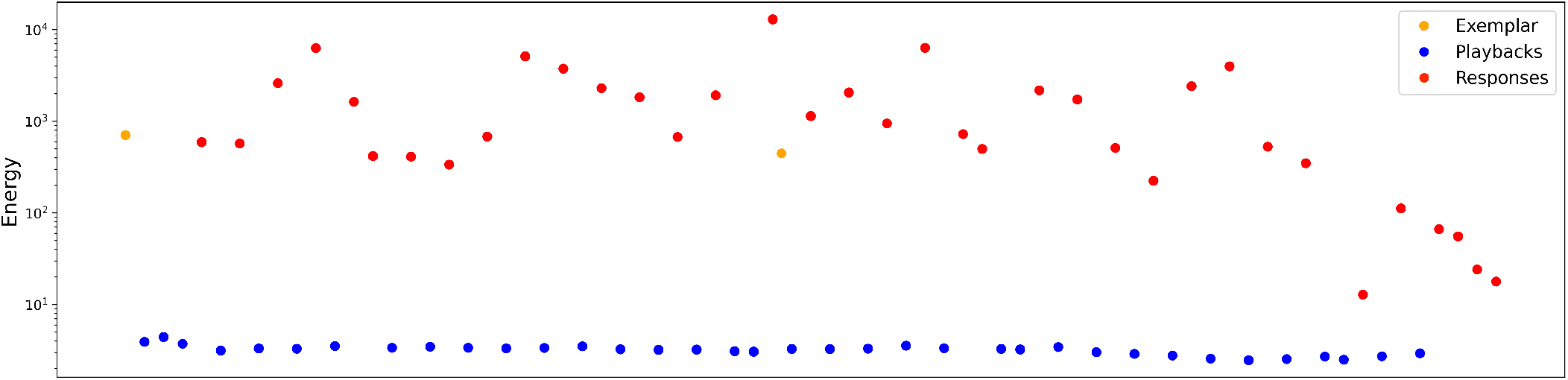
Log plot of the energy of each sound—*P*_0_ *−P*_35_ (blue) and *R*_0_ *−R*_35_ (red) (Left to Right)—in the encounter plus the August 18th exemplar *E*_0_ (orange) (Far Left). Most of Twain’s vocalizations are several orders of magnitude more powerful than the playback sounds produced by the underwater loudspeaker.

### 3. Playback-Response Time Series and Video

These quantitative and visual probes into the sonic signals are all well and good. And, the metrics employed do give novel insight into the encounter. That said, they fall short of measuring semantic content or intention. And so, at the risk of injecting subjectivity, we strongly recommend that the reader simply listen to the full hydrophone recording to come to their own conclusions, especially about communicative aspects not-yet amenable to quantitative analysis.

The hydrophone recording is found at [World Wide Whale. Also](http://worldwidewhale.org/), there is a video recording by top-deck observer (JAH) that captures the surface events during the encounter composed with the hydrophone recording (JPC, BMC) laid over and time aligned.

To appreciate aspects of the nonquantitative communication, when playing either of these the listener should imagine themself in the position of deciding when to broadcast the next playback, having just heard the animal response to the previous.

**Appendix E: Signals and Energy Spectral Density**

The analyzed signals are pulse-like waveforms produced by humpback whales with approximately two second duration, as shown in Fig. S1 (Top, blue). Let *X*(*t*) describe the value of the waveform at time t *∈* (−∞, ∞), where there exists some *k* ∈ *R* such that *X*(*t* > *k*) = 0 and *X*(*t* < − *k*) = 0.

The total energy in each throp is given by:

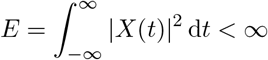

and plotted in Fig. S6.

The Fourier transform of the signal *X*(*t*) at frequency *f* is:

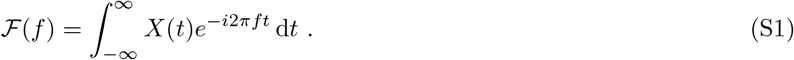

In practice, we use the *discrete-time Fourier transform* (DTFT) given by:

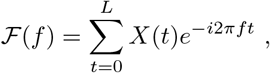

where *L* is the time at which the sample ends.

The *energy spectral density* of *X*(*t*) is defined:

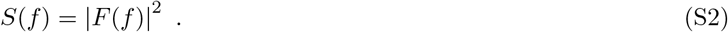

Parseval’s theorem:

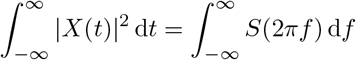

shows that *S*(*f*) represents the energy distribution in the signal as a function of frequency [79, 80].Now, define a finite energy process’ *autocorrelation function* as:

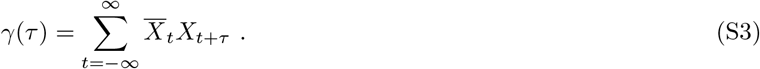

The bar above *X*(*t*) denotes its complex conjugate. Equation (S3) emphasizes that *γ*(*τ*) ignores all of the statistical dependence between *t* and *t* + *τ*. Hence, it is also called the *two-point* correlation function. This makes plain the connection between pairwise statistics and the autocorrelation function.

It is easy to show that:

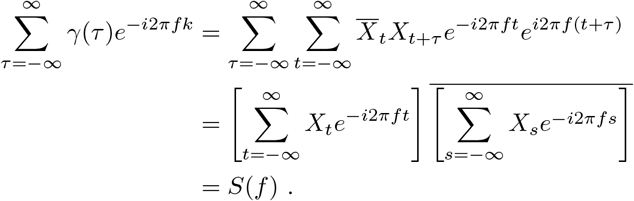

Thus, the energy spectral density can also be obtained as the DTFT of the signal’s autocorrelation *γ*(*τ*).

Directly comparing the energy spectra as shown in Fig. S4 is difficult due to the wide variation in each sound’s total energy, as shown in Fig. S6. To account for this, we normalized the energy spectral density such that:

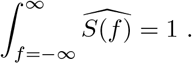

This allows treating each normalized energy spectrum as a probability distribution, where the sum of 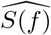over a frequency band indicates the proportion of the the signal’s energy that is delivered within that frequency range.

**Appendix F: Wasserstein Metric**

Figures 7 and 8 compare all of the playback and response normalized energy spectra in terms of their Wasserstein distance. Let’s explain the features of this measure of similarity when applied to energy spectra, the calculation of which is discussed in the prior section (Appendix E).

Let (*M, d*) be a metric space. The Wasserstein distance between two probability measures *µ* and *ν* on *M* is defined [81]:

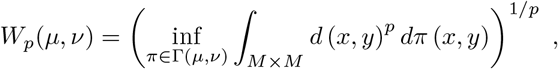

**FIG. S7:**
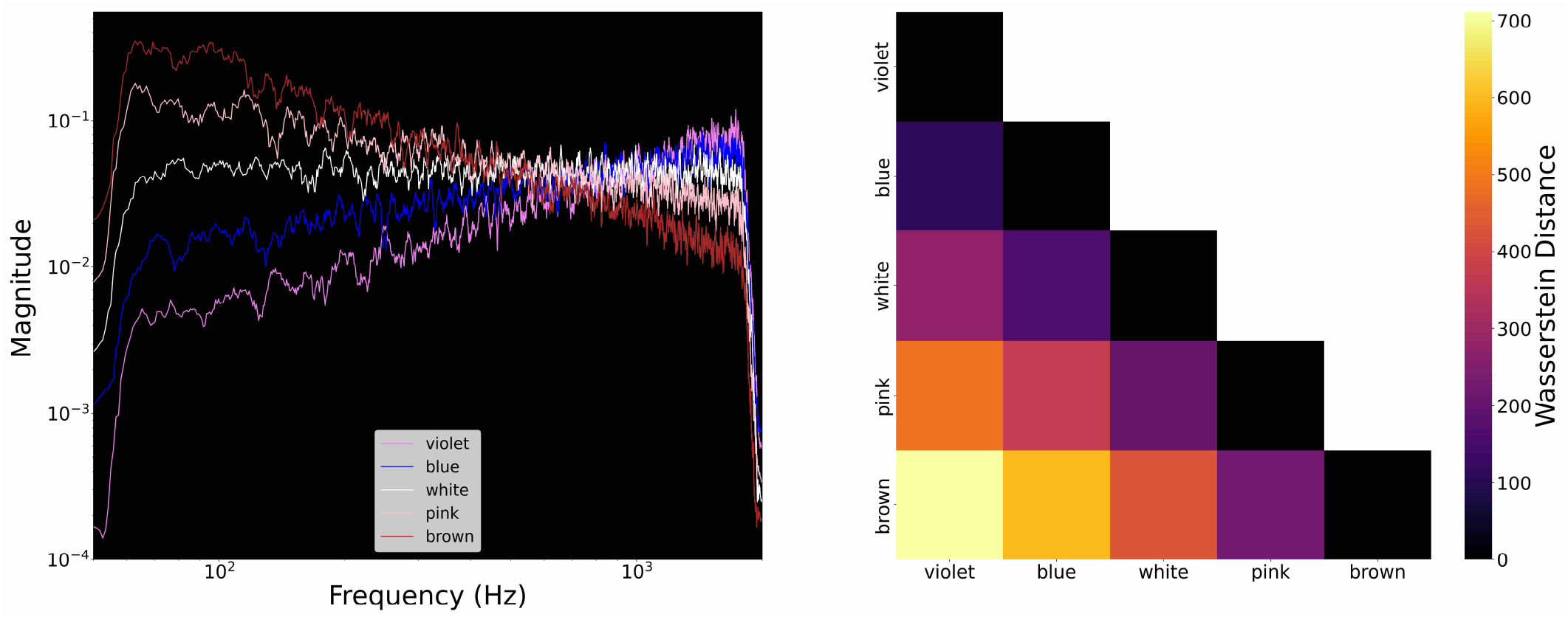
(Left) Log-log plot of the magnitude and frequency of violet, blue, white, pink and brown noise, colored as they are named. (Right) The Wasserstein distance calculated between the different shades of sound between 40 Hz and 2000 Hz.

where Γ(*µ, ν*) is the set of all joint probability distributions that have marginals *µ* and *ν*. It is the minimal cost to shift probability mass from one distribution to match the other’s shape. For *p* = 1, the distance is often called the *earth mover’s distance*.

*W* (*µ, ν*) is the solution to a constrained linear optimization as the objective function and constraints are linear functions of *π*. This is computationally costly, scaling as *O*(*n*^3^ log *n*) in the sample number *n* [82]. However, when *M ⊆* ℝ, there is a closed-form solution to the Wasserstein optimization problem [83]. Let *F* and *G* be the cumulative distributions functions of *µ* and *ν*, respectively. Then:

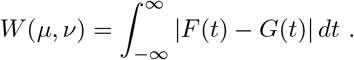

This closed-form solution is considerably faster to compute than the linear optimization required for arbitrary metric spaces.

To compare sounds, recall that Eq. (S2) gives the energy spectral density for finite-energy signals. In effect, we have a library of pulse-like signals that meet this criteria, so we calculate energy spectra using the DTFT. This gives the spectra plotted in Fig. S4. We then normalize the energy spectra such that 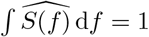. This allows treatingthe energy spectra as probability distributions over frequency and comparing sounds of differing total energy on equal footing.

Take *p* = 1 and the distance metric *d*(·,·) to be the Euclidean metric in frequency space: *d*(*x, y*) = *x y*. Therefore, the Wasserstein distance between any two pure frequencies is simply the difference in their frequencies.

To calibrate intuitions about this metric, we first calculated the Wasserstein distance between the *color noises*. Log-log plots of the energy spectra of the color sounds are given in Fig. S7 (Left). In general the color noises are distinguished by their power spectra. The most well known is white noise, for which all frequencies *f* have the same power: *P* (*f*) = constant. The other color noises are defined by the shape of their energy-frequency curves: pink noise scales as 1*/f*, brown noise 1*/f* ^2^, blue noise as *f*, and violet noise as *f* ^2^.

Note that color noises are generically taken to be infinite random signals and therefore are discussed in terms of their *power* spectral density. In this case, we constructed pulses of approximately two seconds long that demonstrate the color noise energy-frequency curves between 40 Hz to 2000 Hz—the frequency range of the Twain vocalizations. These artificially constructed pulses have finite energy, and so we can treat them like our real world samples.

We take this suite of color pulses and calculate the pairwise Wasserstein distances as shown in Fig. S7 (right). Comparing these results to Fig. 7, we see that the full-spectrum distance between the playbacks and Twain’s responses is roughly equivalent to the distance between pink and blue noise. This makes sense as the responses are weighted in the lower spectral range and the playbacks in the higher, so this is well approximated by comparing 1*/f* to *f*.

We also note that even the closest color noises (blue and violet) are roughly 200 Hz apart—this is on the outer range of distances shown by the comparisons between different Twain responses. That is to say, these sounds are very close indeed. Distinguishing them further requires more sophisticated methods that are beyond the present scope.

One option is to directly compare spectrograms, rather than the energy spectra, which flatten out the time axis. That is to say, taking the DTFT of the signal *X*(*t*) ignores the temporal dynamics of the original signal, indicating only that frequency *f* was present at some time. Applying the Wasserstein metric to compare spectrograms is slightly complicated due to the difficulty of comparing apples to apples, so to speak, with sounds of different lengths, but it is a promising avenue for exploration.

